# Interpreting Temporal Fluctuations in Resting-State Functional Connectivity MRI

**DOI:** 10.1101/135681

**Authors:** Raphaël Liégeois, Timothy O. Laumann, Abraham Z. Snyder, Juan Zhou, B.T. Thomas Yeo

## Abstract

Resting-state functional connectivity is a powerful tool for studying human functional brain networks. Temporal fluctuations in functional connectivity, i.e., *dynamic* functional connectivity (dFC), are thought to reflect dynamic changes in brain organization and *non-stationary* switching of discrete brain states. However, recent studies have suggested that dFC might be attributed to sampling variability of static FC. Despite this controversy, a detailed exposition of stationarity and statistical testing of dFC is lacking in the literature. This article seeks an in-depth exploration of these statistical issues at a level appealing to both neuroscientists and statisticians.

We first review the statistical notion of stationarity, emphasizing its reliance on ensemble statistics. In contrast, all FC measures depend on sample statistics. An important consequence is that the space of stationary signals is much broader than expected, e.g., encompassing hidden markov models (HMM) widely used to extract discrete brain states. In other words, stationarity does not imply the absence of brain states. We then expound the assumptions underlying the statistical testing of dFC. It turns out that the two popular frameworks - phase randomization (PR) and autoregressive randomization (ARR) - generate stationary, linear, Gaussian null data. Therefore, statistical rejection can be due to non-stationarity, nonlinearity and/or non-Gaussianity. For example, the null hypothesis can be rejected for the stationary HMM due to nonlinearity and non-Gaussianity. Finally, we show that a common form of ARR (bivariate ARR) is susceptible to false positives compared with PR and an adapted version of ARR (multivariate ARR).

Application of PR and multivariate ARR to Human Connectome Project data suggests that the stationary, linear, Gaussian null hypothesis cannot be rejected for most participants. However, failure to reject the null hypothesis does not imply that static FC can fully explain dFC. We find that first order AR models explain temporal FC fluctuations significantly better than static FC models. Since first order AR models encode both static FC and one-lag FC, this suggests the presence of dynamical information beyond static FC. Furthermore, even in subjects where the null hypothesis was rejected, AR models explain temporal FC fluctuations significantly better than a popular HMM, suggesting the lack of discrete states (as measured by resting-state fMRI). Overall, our results suggest that AR models are not only useful as a means for generating null data, but may be a powerful tool for exploring the dynamical properties of resting-state fMRI. Finally, we discuss how apparent contradictions in the growing dFC literature might be reconciled.

## 1. Introduction

The human brain exhibits complex spatiotemporal patterns of activity fluctuations even during the resting-state (Greicius et al., 2003; Damoiseaux et al., 2006; Smith et al., 2013b). Characterizing the structure of these fluctuations is commonly done via functional connectivity (FC) analyses of resting-state fMRI (rs-fMRI) data (Van Den Heuvel and Pol, 2010; Buckner et al., 2013). The most common FC measure is the Pearson correlation between brain regional time courses (Biswal et al., 1995; Vincent et al., 2006; Dosenbach et al., 2007; Buckner et al., 2009; Zalesky et al., 2010; Power et al., 2011; Yeo et al., 2011, 2014; Margulies et al., 2016), although other measures, such as partial correlation (Fransson and Marrelec, 2008; Spreng et al., 2013) or mutual information (Tsai et al., 1999; Tedeschi et al., 2005; Chai et al., 2009) have been utilized. These FC measures are *static* in the sense that they are invariant to temporal re-ordering of fMRI time points, thus ignoring temporal information that might be present in fMRI (Theiler et al., 1992; Oppenheim and Willsky, 1997).

In contrast, recent work on *dynamic* functional connectivity (dFC) suggests that there might be important information beyond static FC, e.g., in the temporal fluctuations of FC or in models taking into account the temporal ordering of fMRI time series (see Hutchison et al. (2013a); Calhoun et al. (2014); Preti et al. (2016) for recent reviews). To interrogate dFC, sliding window correlations (SWC) is by far the most common method in human (Sakoğlu et al., 2010; Handwerker et al., 2012; Hutchison et al., 2013b; Allen et al., 2014; Leonardi et al., 2014; Liégeois et al., 2016; Wang et al., 2016) and animal studies (Grandjean et al., 2017), although many alternative approaches have been proposed (Majeed et al., 2011; Smith et al., 2012; Lindquist et al., 2014; Karahanoğlu and Van De Ville, 2015; Shine et al., 2015).

To assess the statistical significance of dFC, randomization frameworks are typically used to generate null data. Null hypothesis testing can then be performed by comparing statistics from the original data against those from the null data. The two most popular randomization frameworks are autoregressive randomization (ARR) (Chang and Glover, 2010; Zalesky et al., 2014) and phase randomization (PR) (Handwerker et al., 2012; Allen et al., 2014; Hindriks et al., 2016). While most papers reported the rejection of the null model (Chang and Glover, 2010; Handwerker et al., 2012; Zalesky et al., 2014), recent studies have suggested difficulties in rejecting the null model, especially in single subject data (Hindriks et al., 2016; Laumann et al., 2016).

The observed dFC has also been interpreted by many authors as evidence of non-stationary switching of discrete brain states (Allen et al., 2014; Hansen et al., 2015). These states have been associated with mental disorders (Damaraju et al., 2014; Rashid et al., 2014; Su et al., 2016; Du et al., 2016), as well as variation in intra-individual and inter-individual differences in vigilance, consciousness and executive function (Barttfeld et al., 2015; Nomi et al., 2017; Shine et al., 2016; Wang et al., 2016). In contrast, some have suggested that the brain (as measured by rsfMRI) might not be undergoing sharp transition between discrete states (Leonardi et al., 2014) or that dFC fluctuations might largely reflect sampling variability (Laumann et al., 2016).

Contributing to the possible confusion in the literature is the loose use of the term “stationarity” (e.g., Hutchison et al., 2013a; Allen et al., 2014; Zalesky and Breakspear, 2015; Preti et al., 2016). For example, Hutchison and colleagues (Hutchison et al., 2013a) equated static FC and dFC analyses with assumptions of stationarity and non-stationarity respectively. However, the very same review cautioned that a stationary process can exhibit temporal fluctuations in an FC metric, such as SWC (Hutchison et al., 2013a). Since null data generation frameworks (e.g., PR and ARR) were developed based on strict statistical definitions of stationarity (Tucker et al., 1984; Efron and Tibshirani, 1986), the loose usage of statistical terminologies impedes our understanding of dFC. To the best of our knowledge, issues of stationarity and assumptions of null data generation frameworks (PR and ARR) are often briefly mentioned, but not discussed in detail in the literature. Exploring these issues in-depth leads to several surprising conclusions.

We begin by clarifying our definitions of several common dFC terms, such as “static”, “dynamics” and “time-varying” (Section 2). A proper explanation of “stationarity” requires more background knowledge. Therefore in the following section, we review random variables, random processes, and weak-sense stationarity, as well as how fMRI can be conceptualized as a random process (Section 3). We then show that a two-state hidden Markov model (HMM; Rabiner, 1989) process is actually stationary, suggesting that stationarity does not necessarily imply the absence of brain states (Section 4). In the following section, assumptions behind PR and ARR are discussed, revealing that both PR and AR generate null data that are linear, stationary and Gaussian. Therefore rejection of the PR and ARR null models does not imply non-stationarity. Importantly, AR models encode dynamical interactions between brain regions, above and beyond static FC (Section 5). Experiments on the Human Connectome Project data suggests that the PR and ARR null models cannot be rejected for most low motion participants, and that bivariate ARR (a common variant of ARR) can yield false positives (Section 6). Furthermore, multivariate AR models replicate the rich dynamics of SWC significantly better than just models of static FC, as well as commonly used HMM-type models that explicitly encode discrete brain states (Section 7). We conclude with a discussion of how these results can be reconciled with the growing literature on dFC (Section 8).

While our experiments focused on SWC, almost all the issues we discuss apply to other dFC methods. In addition, it is worth distinguishing dFC (second order statistics) from dynamic fMRI activity level (first order statistics). From the earliest days of resting-state fMRI, the question of dynamic fMRI activity level during resting-state and its relationship with behavior has been of great interest (Fox et al., 2006, 2007; Kucyi et al., 2016). Many of the issues that we raised in this manuscript also apply to the study of dynamic activity level. Therefore we will point out relevant lessons to dynamic activity level as and when they arise.

## 2. Clarification of “static”, “dynamic” and “time-varying”

In the literature, the terms “static”, “dynamic” and “time-varying” are often not explicitly defined. For example, some authors use the terms “dynamic” and “time-varying” interchangeably. Here we clarify our definitions of these terms, to ensure internal consistency within this manuscript. While we believe our definitions are reasonable, other researchers might prefer different definitions.

First, as mentioned in the introduction, we reserve the term “static” to refer to models or measures that are in-variant to temporal re-ordering of the data points (Theiler et al., 1992). This is in contrast to “dynamic” models or measures that are not invariant to temporal re-ordering. These definitions are consistent with the systems theory literature that have abundantly documented the properties of static (or memoryless) versus dynamical models (e.g., Section 2.2.4 of Theiler et al., 1992; Oppenheim and Willsky, 1997). Finally, “time-varying” measures encode fluctuations over time, while “time-varying” models have parameters that are functions of time (Liégeois, 2015).

**Table 1:** Glossary of different key concepts used in this paper.

| Terms | Explanations | Details |
| --- | --- | --- |
| First order statistics | Statistics based on the random variable raised to the power of 1, e.g., $\mathbb{E}(x)$ or mean, activity level of event-related fMRI responses. | Sec. 1; 3.1 |
| Second order statistics | Statistics based on the random variable raised to the power of 2, e.g., $\mathbb{E}(x^2)$, variance, Pearson’s correlation. | Sec. 1; 3.1 |
| Static | A static measure or model is invariant to temporal re-ordering of the data points, e.g., Pearson’s correlation, mean. | Sec. 2 |
| Dynamic | A dynamic measure or model is affected by temporal re-ordering of the data points, e.g., autocorrelation, lagged covariance, sliding window correlations. A model can be both dynamic and stationary. | Sec. 2 |
| Time-varying | Time-varying measures encode fluctuations over time (e.g., sliding window correlations), while time-varying models have parameters that are functions of time (e.g., GARCH; Bollerslev, 1986). Importantly, stationary processes may exhibit time-varying properties within a single realization. Some dFC literature used the terms “dynamic” and “time-varying” interchangeably. Here we specifically distinguish between the two terms. | Sec. 2 |
| Sample statistics | Statistics computed from a single realization of a random process. Most statistics computed from fMRI time series are sample statistics. | Sec. 3.3 |
| Ensemble statistics | Statistics computed across different realizations of the same random process. We usually don’t have access to ensemble statistics in fMRI. | Sec. 3.3 |
| Stationary | A random process is weak-sense stationary if its first and second order ensemble statistics are constant in time. A stationary process may still exhibit meaningful fluctuations. The space of stationary signals is much larger than one might expect. For example, the hidden Markov model is stationary. | Sec. 3.2 |
| Ergodic | A random process is ergodic if its ensemble statistics are equal to the corresponding sample statistics. | Sec. 3.3 |
| Brain state | In the context of dFC, brain states often refer to distinct FC patterns obtained by clustering SWC time series. There is often an implicit assumption of sharp transition between brain states. | Sec. 4; 8.5 |
| One-lag covariance | The one-lag covariance between two time series is the covariance between the two time series after shifting one of the two time series by one time point. | Sec. 5.1 |
| Autoregressive model | The $p$-th order autoregressive model assumes the signal at time $t$ is a linear combination of the signal at the $p$ previous time points. The first order autoregressive model exactly encodes both static FC and one-lag covariance. | Sec. 5.2 |

For example, Pearson’s correlation of the entire fMRI time courses is static because permuting the ordering of the fMRI frames results in the same values. On the other hand, one-lag covariance (e.g., Eq. (2) in Section 5.1) is dynamic, but not time-varying. 0-th order autoregressive models are static, while higher order autoregressive models are dynamic (see Section 5.2). Similarly, the parameters of the HMM (Rabiner, 1989) and neural mass models (Deco et al., 2011) are not time-varying, although they are both dynamic models. SWC and generalized autoregressive conditional heteroscedastic (GARCH) model (Bollerslev, 1986; Lindquist et al., 2014) are dynamic and time-varying.

It is worth mentioning that the dFC literature does not typically include lagged covariance as examples of dynamic FC and might often equate time-varying FC with dynamic FC. Here we distinguish between dynamic and time-varying because an important result in this article is that first order autoregressive model is able to explain time-varying SWC during resting-state fMRI very well (Section 7), even though first order autoregressive model is only dynamic, but not time-varying. Table 1 summarizes various key terms utilized in this article.

## 3. Interpreting fMRI as a random process

In this section, we first review random variables, random processes and weak-sense stationarity (WSS). We then distinguish between sample statistics and ensemble statistics. Finally, we formalize fMRI as a random process and explain why testing dFC within a formal statistical framework is non-trivial.

### 3.1. Random variables

A random variable is a quantity that is uncertain (Prince, 2012). It may be the outcome of an experiment (e.g., tossing a coin) or real world measurement (e.g., measuring the temperature of a room). If we observe a random variable multiple times, we will get different values. Some values occur more frequently than others; this variation in frequencies is encoded by the probability distribution of the random variable. Multiple observations of a random variable are referred to as *realizations* (or samples) of the random variable.

The mean or expectation of a random variable *X* is denoted as *E*(*X*). We can think of *E*(*X*) as the average value of *X* over many (infinite) realizations of *X*. Similarly, the variance of a random variable *X* is denoted as *V ar*(*X*) = *E*[(*X − E*(*X*))^2^]. We can think of *V ar*(*X*) as the average square deviation of *X* from its mean over many (infinite) realizations of *X*.

In the case of two random variables *X* and *Y*, we can characterize their linear relationship with the covariance *Cov*(*X, Y*) = *E*[(*X − E*(*X*))(*Y − E*(*Y*))]. We can think of *Cov*(*X, Y*) as averaging (*X − E*(*X*))(*Y − E*(*Y*)) over many (infinite) realizations of *X* and *Y*. The covariance measures how much *X* and *Y* co-vary across their respective means. If the covariance is positive, then for a particular realization of *X* and *Y*, if *X* is higher than its mean, then *Y* tends to be higher than its mean. If the covariance is negative, then for a particular realization of *X* and *Y*, if *X* is higher than its mean, then *Y* tends to be lower than its mean. We note that *Cov*(*X, X*) = *V ar*(*X*).

### 3.2. Random processes and weak-sense stationarity (WSS)

A random process is an infinite collection of random variables, and is especially useful for the analysis of time series. For example, suppose we randomly pick a thermometer from a store (with many thermometers) to measure the temperature of a particular room. Let *U_t_* be the thermometer measurement at time *t*. Then *U_t_* is a random process. Figure 1A illustrates three *realizations* of the random process, where each realization (blue, red or green) corresponds to a different thermometer. Here we assume that the room temperature is constant at 20^°^*C*, and that each thermometer is identical and incurs an independent (zero-mean unit-variance Gaussian) measurement noise at each time, i.e., *p*(*U_t_*) *∼ N* (20, 1).

The expectation of a random process *X_t_* at time *t* is denoted as *E*(*X_t_*). We can think of *E*(*X_t_*) as averaging *X_t_* across infinite realizations of the random process at time *t*. For the toy example *U_t_* (Figure 1A), averaging the temperature measurements across many thermometers at a particular time *t* converges to the true temperature of the room, and so *E*(*U_t_*) = 20 for all time *t*. Similarly, we can think of *V ar*(*X_t_*) = *E*[(*X_t_ − E*(*X_t_*))^2^] as averaging the square deviation of *X_t_* at time *t* from its mean *E*(*X_t_*) over infinite realizations of the random process. For the toy example *U_t_* (Figure 1A), *V ar*(*U_t_*) = 1 for all time *t*.

Finally, the auto-covariance *Cov*(*X_n_, X_m_*) = *E*[(*X_n_ − E*(*X_n_*))(*X_m_ − E*(*X_m_*))] measures the co-variation of *X_n_* and *X_m_* about their respective means at times *n* and *m*. For example, if the auto-covariance is positive, then for a particular realization of *X_t_*, if *X_n_* is higher than its mean *E*(*X_n_*), then *X_m_* tends to be higher than its mean *E*(*X_m_*). Conversely, if the auto-covariance is negative, then for a particular realization of *X_t_*, if *X_n_* is higher than its mean *E*(*X_n_*), then *X_m_* tends to be lower than its mean *E*(*X_m_*). We note that *Cov*(*X_n_, X_n_*) = *V ar*(*X_n_*). For the toy example *U_t_* (Figure 1A), *Cov*(*X_n_, X_m_*) = 0 for two different time points *n* and *m* since we assume the thermometer noise is independent at each time point.

We can now define WSS as follows (Papoulis, 2002):

> *A random process X_t_ is WSS if its mean E(X_t_) is constant for all time t and its auto-covariance Cov(X_n_, X_m_) depends only on the time interval Τ = n − m, i.e., Cov(X_n_, X_m_) = R(n − m) = R(Τ)*.

Since *V ar*(*X_n_*) = *Cov*(*X_n_, X_n_*), this implies that *V ar*(*X_n_*) = *R*(0) for a WSS process. In other words, the variance *V ar*(*X_t_*) of a WSS process is constant over time.

While there are other forms of stationarity (e.g., strict-sense stationarity), we will only focus on WSS in this paper and will use the phrase “stationarity” and “WSS” interchangeably. It is also worth mentioning at this point that the dFC community might not be referring to “stationarity” or “non-stationarity” in the strict statistical sense. However, as will be seen in Section 5, the current dFC statistical testing frameworks do rely on the above definition of stationarity. Further discussion is found in Section 8.1.

The toy example *U_t_* (Figure 1A) is WSS because *E*(*U_t_*) = 20 is a constant, and *Cov*(*X_n_, X_m_*) can be written as *R*(*n − m*), where *R*(0) = 1 (when *n* = *m*) and *R*(*n − m*) is equal to 0 for *n − m* ≠ 0. In contrast, suppose the thermostat of the room was changed at time *t*_0_, so that the room temperature increased from 20^°^*C* to 23^°^*C* (Figure 1B). Then the resulting random process *V_t_* is non-stationary because *E*(*V_t_*) = 20 for *t < t*_0_ and *E*(*V_t_*) = 23 for *t > t*_0_.

### 3.3. Ensemble statistics versus sample statistics

It is worth emphasizing that the mean *E*(*X_t_*), variance *V ar*(*X_t_*) and auto-covariance *Cov*(*X_n_, X_m_*) of a random process are defined *across* an infinite number of realizations, rather than within a *single* realization. To illustrate this point, suppose at time points *t*_1_ and *t*_2_, we average across multiple realizations of the random process *U_t_* resulting in *M_U_* (*t*_1_) and *M_U_* (*t*_2_) (Figure 1A). We can think of *M_U_* (*t*_1_) and *M_U_* (*t*_2_) as estimates of *E*(*U_t_*) and *E*(*U_t_*_2_ Indeed, as the number of realizations increases, *M_U_* (*t*_1_) and *M_U_* (*t*_2_) will converge to *E*(*U_t_*_1_) and *E*(*U_t_*_2_) respectively. This convergence holds not just for WSS processes, but all random processes. In the toy example *V_t_* (Figure 1B), *M_V_* (*t*_1_) converges to *E*(*V_t_*_1_) = 20, while *M_V_* (*t*_2_) converges to *E*(*V_t_*_2_) = 23. We refer to the computation of statistics across realizations as *ensemble* statistics.

**Figure 1:**
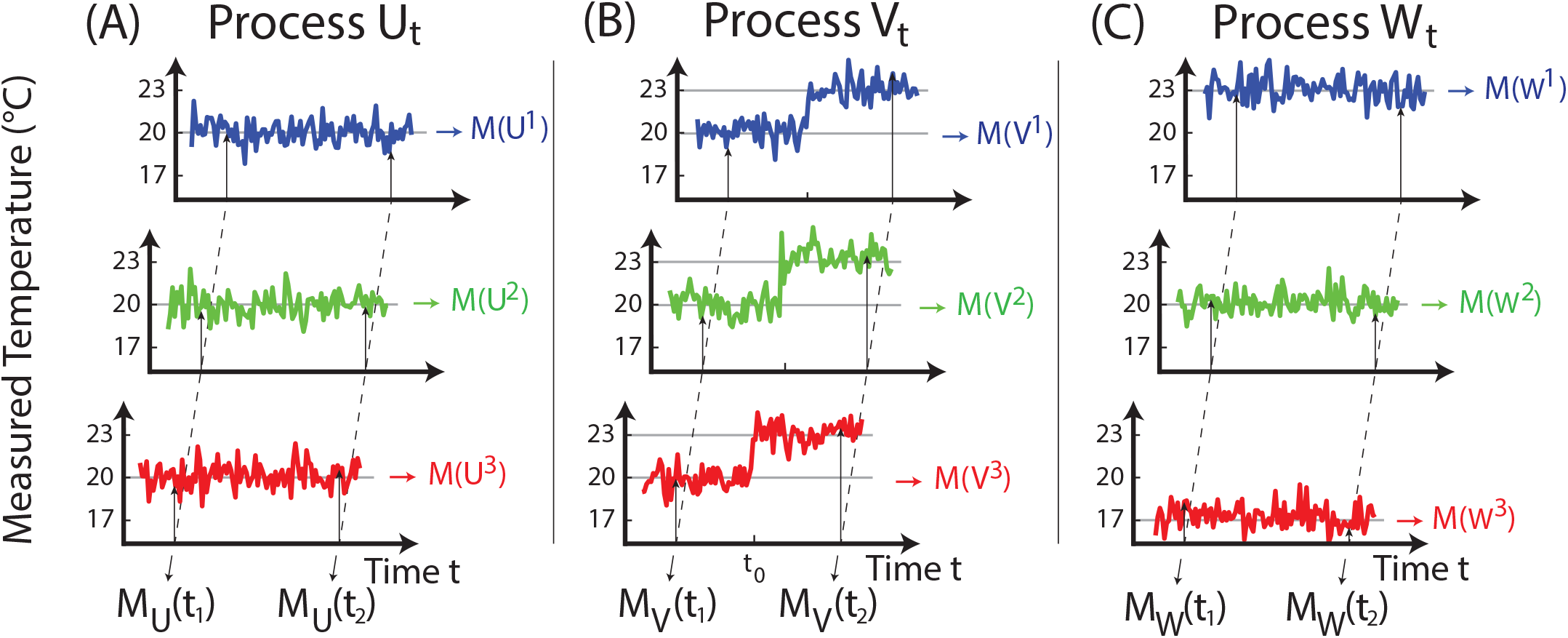
Three examples of random processes *U_t_* (left), *V_t_*(middle) and *W_t_*(right) and three realizations in each case illustrating the distinction between ensemble and sample statistics. Ensemble statistics are computed across realizations, while sample statistics are computed within a single realization. The random process *U_t_*is weak-sense stationary (WSS) because ensemble statistics (e.g., mean and variance) do not depend on time, i.e., *M_U_* (*t*_1_) = *M_U_* (*t*_2_). *U_t_* is also ergodic because ensemble statistics are equal to sample statistics, i.e., *M_U_* (*t*_1_) = *M* (*U* ^1^) = *M* (*U* ^2^) = *M* (*U* ^3^). Random process *V_t_* is not WSS because the ensemble mean is not constant in time, i.e., *M_V_* (*t*_1_) ≠ *M_V_* (*t*_2_). It is also not ergodic, i.e., *M_V_* (*t*_1_) *M* (*V* ^1^). Random process *W_t_* is WSS, i.e., *M_W_* (*t*_1_) = *M_W_* (*t*_2_). However, *Wt* is not ergodic because ensemble statistics are constant in time but sample mean is not the same for different realizations, i.e., *M_W_* (*t*_1_) */*≠*M* (*W* ^1^) */*≠*M* (*W* ^2^)≠ *M* (*W* ^3^).

In contrast, *sample* statistics are computed within a single realization. For example, we can average each realization of the random process *U_t_* resulting in *M* (*U* ^1^), *M* (*U* ^2^) and *M* (*U* ^3^) (blue, green and red in Figure 1A). In the case of the random process *U_t_* (Figure 1A), as the number of time points for each realization increases, the sample statistics converge to the ensemble statistics. More specifically, *M* (*U* ^1^), *M* (*U* ^2^) and *M* (*U* ^3^) converge to *E*(*U_t_*) = 20. However, sample statistics do not converge to ensemble statistics for non-stationary processes. In the toy example *V_t_* (Figure 1B), the sample statistics *M* (*V* ^1^), *M* (*V* ^2^) and *M* (*V* ^3^) (blue, green and red in Figure 1B) converge to 21.5.

Therefore ensemble statistics are generally not equivalent to sample statistics in the case of non-stationary processes. Based on the toy example *U_t_* (Figure 1A), one might be tempted to conclude that ensemble and sample statistics are equivalent in WSS processes. However, this turns out not to be true. To illustrate this, let’s again assume the room temperature is constant at 20^°^ *C*. However, each thermometer now incurs an independent nonzero-mean unit-variance Gaussian noise at each time *p*(*W_t_*) ~*N* (20 + *b*, 1), where the bias *b* ~ *N* (0, 1) is different for each thermometer (but held constant within a realization). Three realizations of random process *W_t_* are illustrated in Figure 1C. The random process *W_t_* is WSS with *E*(*W_t_*) = 20 (because the bias *b* is zero-mean), *V ar*(*W_t_*) is constant over time, and *Cov*(*W_n_, W_m_*) = 0 for *n /*= *m*. The ensemble mean *M_W_* (*t*_1_) and *M_W_* (*t*_2_) still converge to *E*(*W_t_*) = 20. However, the sample means *M* (*W* ^1^), *M* (*W* ^2^) and *M* (*W* ^3^) now converge to 17, 20 and 23 respectively because we have assumed *b* = – 3, 0, 3 for the blue, green and red realizations in Figure 1C. We now define ergodicity as follows (Papoulis, 2002):

> *A random process X_t_ is ergodic if its ensemble statistics and sample statistics converge to the same values. An ergodic process is WSS*.

The distinction between ensemble statistics and sample statistics becomes important as we conceptualize fMRI as a random process in the next section.

### 3.4. Interpreting fMRI as a random process

In the previous examples of random processes (Figure 1), each realization consisted of one uncertain quantity (temperature) at each time point. However, a random process *X_t_* can also be *multivariate*, i.e., each realization consists of a vector at each time point. fMRI data typically consists of *N* time series, where *N* is the number of voxels or regions of interest (ROIs). Therefore the fMRI data can be thought of as a multivariate random process with an *N* × 1 vector of measurements at each time point (i.e., TR).

For a multivariate random process, *E*(*X_t_*) is now an *N ×*1 vector equivalent to averaging *X_t_* across infinite realizations of the random process at time *t*. The *N × N* auto-covariance matrix *Cov*(*X_n_, X_m_*) = *E*[(*X_n_ − E*(*X_n_*))(*X_m_*−*E*(*X_m_*))^*T*^] measures the co-variation of *X_n_* and *X_m_* about their respective vectorial means at times *n* and *m*. For example, if the *i*-th row and *j*-th column of the auto-covariance matrix is positive, then for a particular realization of *X_t_*, if the *i*-th element of *X_n_* is higher than its mean (the *i*-th element of *E*(*X_n_*)), then the *j*-th element of *X_m_* tends to be higher than its mean (the *j*-th element of *E*(*X_m_*)). For a WSS process, the covariance *Cov*(*X_n_, X_n_*) = *E*[(*X_n_ − E*(*X_n_*))(*X_n_ − E*(*X_n_*))^*T*^] is constant over time.

In the case of fMRI, one could potentially interpret *Cov*(*X_n_ , X_n_*) as the *ensemble N ×N* (un-normalized) functional connectivity matrix among all brain regions at time *n* and might therefore be relevant for the topic of dFC. However, difficulties arise because the auto-covariance and WSS are based on ensemble statistics, and therefore require multiple realizations of a random process to estimate. While most researchers can probably agree that the fMRI data of a single subject can be considered a single realization of a random process, what constitutes multiple realizations is more ambiguous. Most neuroscientists would probably balk at conceptualizing the fMRI data of each subject (of a multi-subject dataset) as a single realization of the *same* random process. Therefore in most dFC papers (Chang and Glover, 2010; Handwerker et al., 2012; Zalesky et al., 2014; Hindriks et al., 2016), the fMRI data of different subjects are treated as single realizations of *different* random processes^1^. As such, both ensemble statistics and hypothesis testing only have access to a single realization of a random process (i.e., relying on sample statistics). Yet, for sample statistics to converge to ensemble statistics (previous section), fMRI must be ergodic, which in turn implies WSS. This creates conceptual and practical issues for dFC analyses that will be the focus for the remainder of this paper.

## 4. Stationarity does not imply the absence of brain states

The previous section suggests the existence of conceptual and practical issues when studying dFC. This section focuses on the conceptual issue of whether non-WSS and dFC are equivalent. In the literature, it is often implicitly assumed that WSS implies the lack of fluctuations in FC (e.g., as measured by SWC) or FC states (e.g., Allen et al., 2014). However, we now show that WSS does not imply the lack of FC fluctuations or FC states.

Consider a toy "brain" with two regions whose signals correspond to a bivariate random process X_t_ containing two brain states *S*_1_ and S_2_. If the brain is in state S1 at time *t*, then 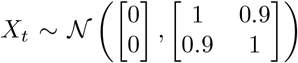,i.e., the two brain regions are functionally connected with *r* = 0.9. If the brain is in state *S*_2_ at time *t*, then X_t_ ~ 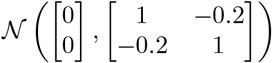 i.e., the two regions are anticorrelated with r = –0.2. Finally, let the probability of transitioning between the two brain states be given by the following transition probability matrix: 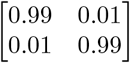, i.e., from time *t* to *t* + 1, there is a 0.99 probability of remaining in the same state and a 0.01 probability of switching state. This random process is known as a hidden Markov process (HMM) (Baum and Petrie, 1966).

**Figure 2:**
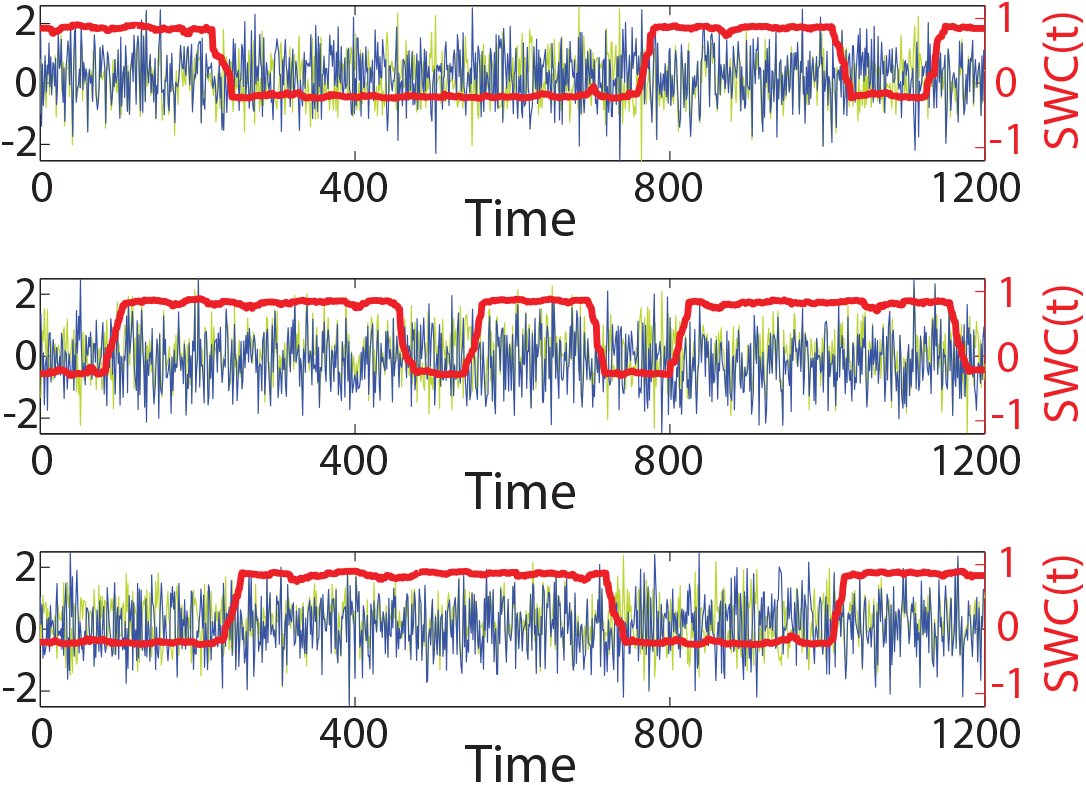
Three realizations of the Hidden Markov Model (HMM) process with two brain regions and two states. Time series of the two brain regions are shown in blue and light green (scale on the left vertical axis). The sliding window correlation (SWC) between the two time series (number of frames *T* = 1200, window size *w* = 30 frames) is shown in red (scale on the right vertical axis). The HMM process is stationary, yet exhibits abrupt transitions in SWC corresponding to the switching between brain states.

Three realizations of this process are shown in Figure 2. The blue and green time courses correspond to the signals of the two brain regions. The sliding window correlations (SWC) between the two regions (red lines in Figure 2) exhibit huge fluctuations with correlations close to 0.9 and –0.2 in states *S*_1_ and *S*_2_ respectively.

Most neuroscientists would probably agree that this toy brain exhibits brain states and dFC. However, this toy brain is also WSS because the ensemble mean *E*(*X_t_*) is constant over time, and the ensemble auto-covariance *Cov*(*X_n_, X_m_*) is only a function of the interval *n – m*. More importantly, the ensemble (unnormalized) functional connectivity matrix *Cov*(*X_n_, X_n_*) in this toy brain is constant over time, despite the presence of brain states and dFC (Figure 2). This situation arises because dFC and brain states exist within a single realization, where the FC at a given time depends on the underlying brain state, which can change over time in a single realization. However, these dynamics are averaged out (across realizations) when considering ensemble notions like WSS. Therefore WSS does not imply the absence of brain states or fluctuations in FC^2^.

It is also worth noting that non-WSS does not necessarily imply the presence of dFC either. If the (unnormalized) functional connectivity matrix *Cov*(*X_n_, X_n_*) varies as a function of time, then fMRI is non-WSS and the brain exhibits true fluctuations in FC. However, non-WSS can arise from just a non-stationary mean *E*(*X_t_*). In other words, the first order statistics (mean) can be non-stationary, while the second order statistics (variance and covariance) remains stationary. For example, the random process *V_t_* (Figure 1A) is non-stationary because its mean *E*(*V_t_*) is non-constant over time, but its variance *V ar*(*V_t_*) is actually constant over time. Therefore, even if fMRI is shown to be non-WSS, this might be due to non-stationary spontaneous activity level and/or non-stationary FC. In other words, non-WSS does not necessarily imply real fluctuations in FC.

While we focus on dFC (second order statistics) in this paper, many of the same issues also apply to the study of dynamic activity level (first order statistics). For example, one could modify the previous HMM example (Figure 2) so that the two states have different ensemble means, but the same ensemble covariance matrix. In this case, the resulting toy brain will be WSS, while still exhibiting real dynamic spontaneous activity (but not dFC)^3^.

Despite the caveat that non-WSS does not necessarily imply real fluctuations in FC, establishing non-stationarity of fMRI is still a useful step towards establishing dFC. Therefore in the next section, we leave behind the conceptual issues raised in this section, and dive into the statistical testing of non-stationarity when only a single realization of the random process is available.

## 5. Stationarity cannot be tested alone

Statistical testing of FC non-stationarity is difficult for two reasons. First, observed dFC values (e.g., using SWC) are only estimates of true values (Hindriks et al., 2016; Laumann et al., 2016). As such, observed dFC fluctuations might simply correspond to sampling variability or measurement noise. Second, as explained in previous sections, the notion of stationarity is based on ensemble statistics defined across infinite realizations of a random process. Therefore observing fluctuations in a single realization of fMRI cannot be directly interpreted as evidence for non-stationarity but requires further statistical testing.

This section seeks to provide insights into common approaches for statistical testing of FC stationarity. A statistical test for non-stationarity requires defining a test statistic and a procedure to generate null data preserving certain properties of the single fMRI realization. The test statistic computed from real data is then compared against the null distribution of test statistics computed from the null data. A significant deviation of the real test statistic from the null distribution of test statistics would result in the rejection of the null hypothesis.

The test statistic should ideally reflect the null hypothesis being tested. For example, if one is interested in whether there are “real” fluctuations in FC between two brain regions, then an intuitive test statistic might be the variance of the SWC (Hindriks et al., 2016):

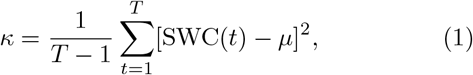

where SWC(*t*) is the SWC between the two brain regions at time *t* and *µ* is the mean of the SWC time series. A higher *κ* (relative to a properly generated null distribution) indicates stronger evidence of dFC.

While there have been a wide variety of test statistics and dFC measures proposed in the literature (Sakoğglu et al., 2010; Chang and Glover, 2010; Zalesky et al., 2014; Hindriks et al., 2016), there has been significantly less discussion about the assumptions behind procedures for generating null data. Such assumptions are highly important because if one manages to generate WSS null data that also preserve all other properties (e.g., possible nonlinearity or non-Gaussianity) of the original data, then a rejection of the null hypothesis would imply non-WSS. However, as will be seen, the two main approaches – autoregressive randomization (ARR) and phase randomization (PR) – actually generate linear, WSS, Gaussian data. Therefore a rejection of the null hypothesis only implies that the signal is nonlinear or non-WSS or non-Gaussian or any combination of the above (Schreiber and Schmitz, 2000).

We begin by discussing properties of the fMRI time series we hope to preserve in the null data, followed by explaining how ARR and PR preserve these properties, and the relationships between the two approaches. Finally, we illustrate with examples of how the null hypothesis can be rejected even though the underlying data is WSS.

### 5.1. Properties to preserve in null data

To test if observed fluctuations in FC (e.g., SWC) can be completely explained by static FC (e.g., Pearson correlation), the null data should retain the static FC observed in real data. To preserve static FC, we could simply generate null data by permuting the temporal ordering of fMRI time courses. However, this procedure destroys the auto-correlational structure inherent in fMRI, and therefore the null hypothesis will be easily rejected. In other words, procedures for generating null data should also preserve the auto-correlational structure of fMRI data in addition to static FC.

Let us define these auto-correlations more precisely. Suppose we observe fMRI time courses from *N* ROIs (or voxels) of length *T*. Let *x_t_* be the *N* × 1 vector of fMRI data at time *t* after each time course has been demeaned. We define the auto-covariance sequence to be the following *N × N* matrices:

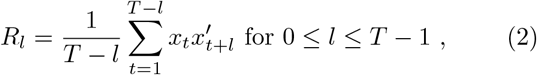

Where *’* denotes transpose. We note that the diagonal elements of *R_l_* encode the *auto*-covariance of individual time courses, while the off-diagonal terms of *R_l_* encode what is usually referred to as the *cross*-covariance between pairs of time courses.

The auto-covariance sequence {*R_l_*} measures the co-variance of fMRI data *l* time points apart. Since the auto-covariances are computed from a single realization of fMRI data, the *R_l_* are considered sample statistics. *R*_0_ is the (un-normalized) functional connectivity matrix typically computed in the literature. If fMRI data is ergodic (and hence WSS), then *R*_0_ would be equal to the *ensemble* (un-normalized) functional connectivity matrix among all brain regions (for sufficiently large *T*). As will be seen, *R*_0_ encodes static properties of fMRI, while the higher order auto-covariances *R*_1_,*…, R_T −_*_1_ encode the dynamic properties of fMRI. We will now examine two frameworks for generating null data that preserve auto-covariances of the original data.

### 5.2. Autoregressive randomization (ARR)

The ARR framework has been utilized by the statistics and physics community for decades (Efron and Tibshirani, 1986) and adopted by seminal papers in the dFC literature (Chang and Glover, 2010; Zalesky et al., 2014). Suppose we have fMRI time courses from *N* brain regions (or voxels). Each fMRI time course is assumed to be demeaned^4^. ARR assumes that the fMRI data at time *t* is a linear combination of the fMRI data from the previous *p* time points:

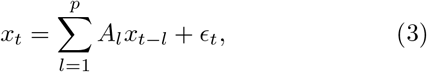

where *p ≥* 1, *x_t_* is the *N ×* 1 vector of fMRI data at time *t*, *∈*_*t*_ ~*N* (0, Σ) corresponds to independent zero-mean Gaussian noise^5^, and *A_l_* is an *N × N* matrix encoding the linear dependencies between time *t* and time *t −l*. Eq. (3) is known as a *p*-th order Gaussian autoregressive (AR) model.

ARR proceeds by first estimating the AR model parameters (Σ, *A*_1_,*…, A_p_*) from the fMRI data (details in Appendix A1). Each null fMRI time series is initialized by randomly selecting *p* consecutive time points of the original data, and then repeatedly applying the AR model (Eq. (3)) to generate *T – p* new time points until null data of length *T* are generated.

> *Suppose the estimated AR model is stable. Then the AR model corresponds to a linear WSS Gaussian process whose auto-covariance sequence R_0_,… R_p_ matches those of the original data*.

The preservation of the first *p*+1 auto-covariances of the original data is a consequence of the Yule-Walker equations (Yule, 1927; Walker, 1931). Further details are found in Appendix A2. One consequence of the above result is the need to verify that the estimated AR model parameters correspond to a stable AR model (see Appendix A2).

Furthermore, any Gaussian linear process can be approximated arbitrarily well by an AR model^6^ (El-Shaarawi and Piegorsch, 2013). Therefore, significant deviation from ARR null data (i.e., null hypothesis rejected) might be due to the fMRI data being nonlinear, non-Gaussian, non-WSS or any of the above.

The matching of the higher order auto-covariances *R*_1_, *…, R_p_* arise from the linear dynamical interactions between brain regions (*A_l_* in Eq. (3)). Therefore the higher order auto-covariances encode the dynamic properties of functional connectivity beyond the static FC encoded by *R*_0_.

### 5.3 Phase randomization (PR)

The PR framework for generating null data has been utilized in the physics community for decades (Tucker et al., 1984; Osborne et al., 1986; Theiler et al., 1992; Prichard and Theiler, 1994). It has been applied to fMRI in several important dFC papers (Allen et al., 2014; Hindriks et al., 2016). We again assume without loss of generality that each fMRI time courses has been demeaned. The PR procedure generates null data by performing Discrete Fourier Transform (DFT) of each time course, adding a uniformly distributed random phase to each frequency, and then performing the inverse DFT. Importantly, the random phases are generated independently for each frequency, but are the *same* across brain regions^7^.

> *PR generates data corresponding to a linear WSS Gaussian process whose auto-covariance sequence R_0_,…, R_T −1_ matches those of the original data*.

Proof of WSS is found in Appendix A4. The preservation of all auto-covariances of the original data is a consequence of the Wiener-Khintchine theorem (Wiener, 1930; Khintchine, 1934; Prichard and Theiler, 1994; Weisstein, 2016); see further elaborations in Appendix A5. An important consequence of the above result is that a rejection of the null hypothesis could be due to the fMRI data being non-linear, non-Gaussian, non-WSS or any of the above.

The relationship between ARR and PR is summarized in Figure 3. ARR preserves the first *p* +1 auto-covariances of the original data, while PR preserves the entire auto-covariance sequence. Therefore, if the original data is not auto-correlated beyond *p* time points (i.e., *R_l_* = 0 for *l* greater than *p*), then a *p*-th order ARR would be theoretically equivalent to PR, except for implementation details. Differences arising from implementation details should not be downplayed. For example, estimating the parameters of a *p*-th order AR model requires the original data to be at least of length *T* = *p* (*N* + 1), where *N* is the number of brain regions. The implication is that a (*T* – 1)-th order ARR cannot be performed even if it is theoretically equivalent to PR. Another difference is that PR null data is only Gaussian for sufficiently long *T* (Tucker et al., 1984), while ARR does not have the constraint. On the other hand, ARR preserves the auto-covariance sequence for sufficiently long *T*, while PR preserves the entire auto-covariance sequence for any *T*. Yet another difference is that PR can only generate null data of the same length as the original data, while ARR can generate null data of arbitrary length, although in the case of dFC, we are typically interested in generating null data of the same length as the original data. As will be seen in the next sections, both approaches appear to yield similar conclusions in dFC analyses despite the practical differences.

**Figure 3:**
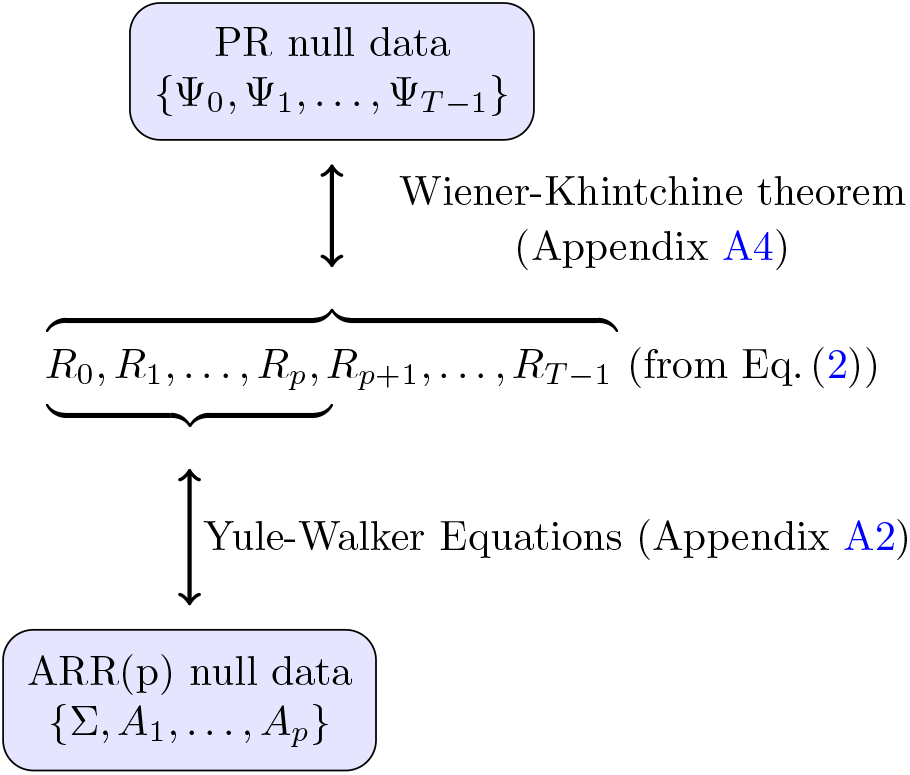
Properties of original data preserved in autoregressive randomized (ARR) and phase randomized (PR) null data. By preserving the power spectral density of the original data, the Wiener-Khintchine theorem (Appendix A4) ensures that PR null data preserve the full original auto-covariance sequence *R*0, *R*1, *…, R_T −_*_1_ (Eq. (2)), where *T* is the number of time points. On the other hand, ARR null data generated from an AR(p) model (Eq. (3)) preserve the first *p* + 1 terms of the auto-covariance sequence via the Yule-Walker equations (Appendix A2).

### 5.4. Stationary but nonlinear or non-Gaussian data can be rejected by ARR and PR

To demonstrate that rejection of the null hypothesis with ARR and PR null data does not imply non-stationarity, we consider the toy brain in the previous section (Figure 2). Recall that the toy brain is a HMM with two brain regions and two brain states, but is WSS.

Figure 4 shows a single realization of this toy brain (Figure 4A), and corresponding first order ARR (Figure 4B) and PR (Figure 4C) null data. The blue and green time courses correspond to the signals of the two brain regions. The PR and ARR null data successfully replicate the auto-covariance sequence of the original time series. For example, the static functional connectivity (Pearson correlation) between the time courses of the two brain regions is equal to 0.35 in the original, ARR and PR data.

**Figure 4:**
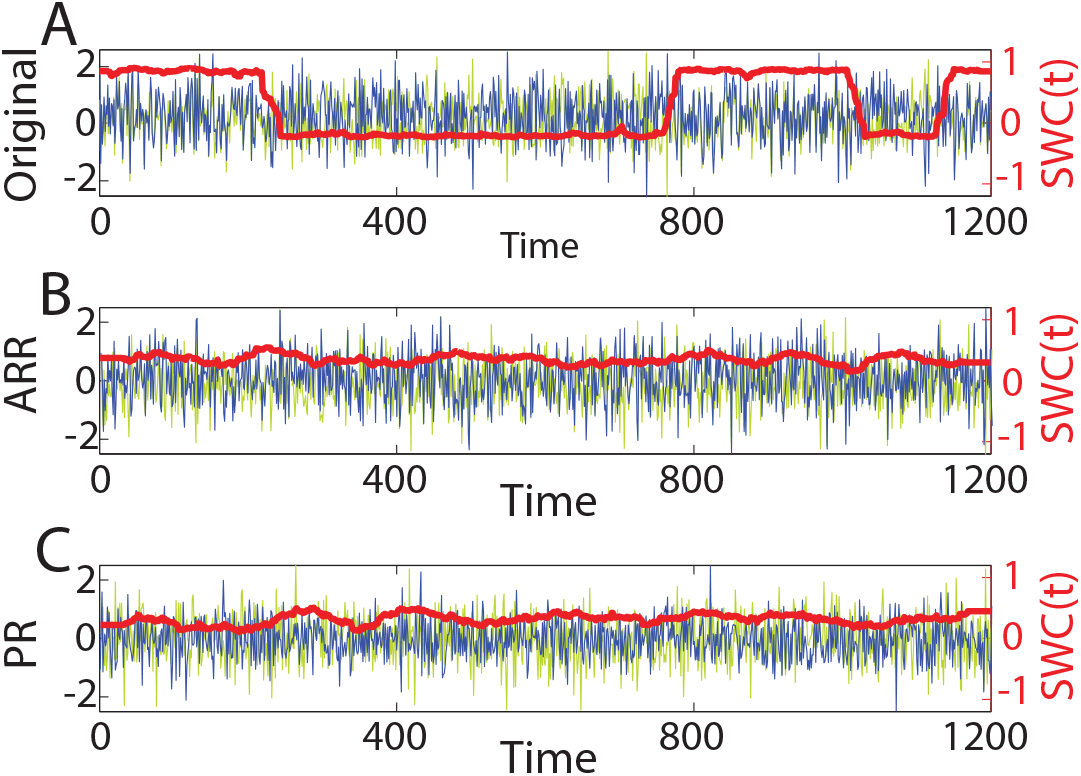
Examples of ARR and PR null time series generated from one realization of the two-state Hidden Markov Model (HMM) process with two brain regions. Time series of the two brain regions are shown in blue and light green (scale on the left vertical axis). The sliding window correlation (SWC) between the two time series (number of frames *T* = 1200, window size *w* = 30 frames) is shown in red (scale on the right vertical axis). (A) The original data exhibits sharp transitions in SWC. (B, C) The null data do not exhibit sharp transitions in SWC.

On the other hand, the SWC between the two regions in the original data (red line in Figure 4A) exhibit huge fluctuations with correlations close to 0.9 and − 0.2 in states *S*1 and *S*2 respectively. However, there is little variation in SWC correlations for the null data (red lines in Figures 4A and 4B). Using the *κ* statistic (Eq. (1)), the null hypothesis is easily rejected.

This result is indeed good news because the implication is that using current methodologies, the null hypothesis can be rejected for a WSS brain with states assuming sufficient statistical power (e.g., the states have sufficiently distinct connectivity patterns and the sliding window is shorter than the average dwell time of a brain state, etc). However, the bad news is that a rejection of the null hypothesis does not imply the existence of brain states because the rejection might simply be due to non-Gaussianity. For example, Figure S1 illustrates a linear, WSS, non-Gaussian process (with no latent states), where the stationarity, linear, Gaussian null hypothesis is easily rejected.

In the event that the stationary, linear, Gaussian null hypothesis is rejected, more advanced approaches can be utilized to differentiate the underlying causes. The amplitude adjusted PR (AAPR; Theiler et al., 1992) controls for non-Gaussianity in the original data by generating linear, stationary data whose amplitude distribution matches the original data. For example, the AAPR null hypothesis is not rejected for the previous linear, WSS, non-Gaussian process example (Figure S1), thus indicating that the rejection of the PR null hypothesis is simply due to non-Gaussianity. Nevertheless, these more advanced considerations are moot because experiments with real data (next section) suggest that the stationary linear Gaussian model cannot be rejected for most low motion Human Connectome Project (HCP) participants.

## 6. stationary linear Gaussian model cannot be rejected for most low motion subjects

In this section, we show that for most low-motion Human Connectome Project (HCP) participants, the stationary linear Gaussian model cannot be rejected. In addition, we show that one form of ARR used in the literature might result in false positives and should be utilized with care.

### 6.1. HCP data and SWC computation

We considered ICA-FIX fMRI data from the HCP S900 data release in fsLR surface space (Glasser et al., 2013; Smith et al., 2013a; Van Essen et al., 2013). Since motion can potentially introduce false positives in dFC (Laumann et al., 2016), our analyses were restricted to participants whose maximum framewise root mean square (FRMS) motion^8^ was less than 0.2mm and maximum DVARS was smaller than 75. Among the four fMRI runs available for each HCP participant, the second run (REST1 RL) yielded the most participants (116) who survived these criteria and was therefore considered (the remaining runs were ignored). Of the 116 remaining participants (or runs), the top 100 participants with the smallest average FRMS were selected. Among these 100 low motion participants, average FRMS of the second run ranged from 0.051mm to0.073mm.

For each participant, the fMRI signal was averaged within each of 114 cortical ROIs (Yeo et al., 2011, 2015; Krienen et al., 2016) resulting in an 114 × 1200 matrix of fMRI data per participant. Following Zalesky et al. (2014), SWC was computed using a window size of 83 frames (59.76s), consistent with window sizes recommended in the literature (Leonardi and Van De Ville, 2015; Liégeois et al., 2016). There was no temporal filtering, except for the very gentle highpass filtering (2000s cutoff) applied by the HCP team (Smith et al., 2013a).

### 6.2. PR, multivariate ARR and bivariate ARR

For each participant, null fMRI data were generated using ARR (section 5.2) and PR (section 5.3). For the ARR procedure, the most common variant in the literature (Chang and Glover, 2010; Zalesky et al., 2014) in^-^ volves estimating for each pair of brain regions, a 2 *×* 2 *A_l_* matrix (Eq. (3)) for each temporal lag *l* (even though there are 114 ROIs). In other words, the resulting null time courses are generated for each pair of brain regions separately. We refer to this procedure as bivariate ARR. In contrast, multivariate ARR estimates a single 114 × 114 *A_l_* matrix (Eq. (3)) for each lag *l*. For multivariate ARR, an AR order of *p* = 1 was utilized. For bivariate ARR, an AR order of *p* = 11 was utilized (Zalesky et al., 2014). Changing the order *p* did not significantly affect our conclusions (see additional control analyses in section 6.6). For each procedure and each participant, 1999 null datasets were generated^9^.

### 6.3 SWC of most pairs of brain regions exhibit stationary, linear and Gaussian dynamics

We first tested if there exists “dynamic” connections in the human brain, defined as ROI pairs exhibiting greater SWC variance (Eq. (1)) than those from null data. To this end, for each participant and ROI pair, the observed SWC variance (computed from real data) was compared against the null distribution of SWC variance generated from the 1999 null datasets, resulting in one *p* value for each ROI pair^10^. Within each participant, multiple comparisons were corrected by applying a false discovery rate (FDR) of *q <* 0.05 to the 6441 *p* values.

Figure 5 illustrates the number of significant ROI pairs across the 100 participants. For both multivariate ARR and PR, 57% of the participants have 0 significant edges. On average (across 100 participants), 36.8 and 34.2 edges were significant with multivariate ARR and PR respectively. Therefore a stationary linear Gaussian model was able to reproduce the SWC fluctuations of more than 99.4% of ROI pairs.

**Figure 5:**
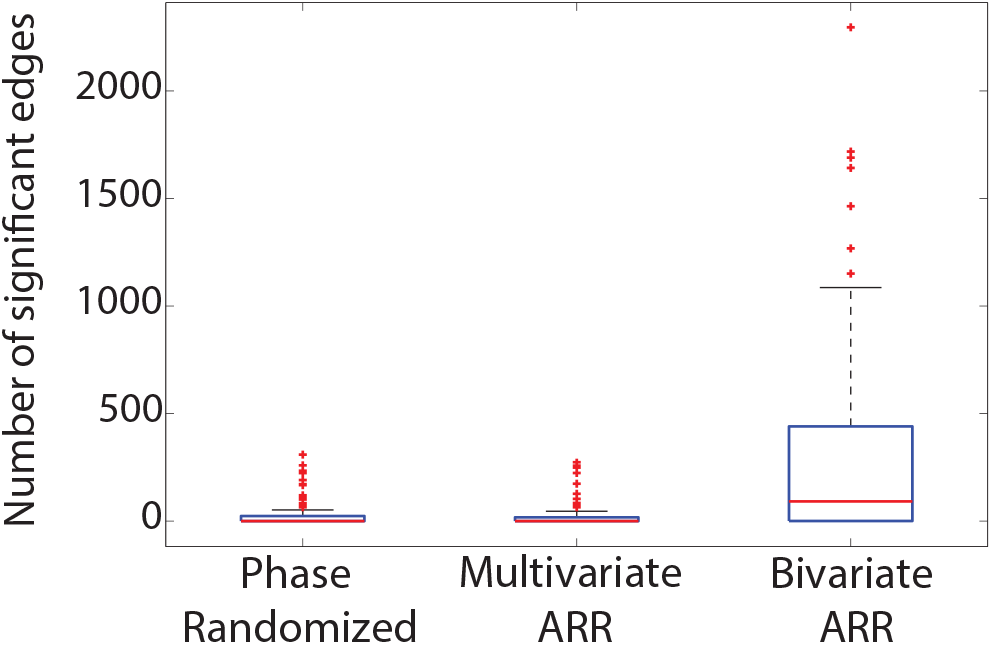
Number of edges in each participant exhibiting significantly greater SWC variance than (i) phase randomized (PR) null data, (ii) multivariate ARR null data and (iii) bivariate ARR null data. On average (across 100 participants), less than 40 (out of 6441) edges were statistically significant for PR and multivariate ARR. For bivariate ARR, 306.7 edges were statistically significant on average (across 100 participants).

On the other hand, for bivariate ARR, 21% of the participants have 0 significant edges. On average (across 100 participants), 306.7 edges were significant. In other words, bivariate ARR tends to be less strict in terms of rejecting the null hypothesis. We will return to this point in Section 6.5.

### 6.4. For almost all low-motion HCP subjects, coherent brain dynamics are stationary, linear and Gaussian

Existence of *coherent* SWC fluctuations was tested using the approach of the pioneering dFC paper (Zalesky et al., 2014). More specifically, this approach involves computing the SWC time series for all ROI pairs and then selecting the top 100 most dynamic SWC time series as measured by the SWC variance (Eq. (1)). The percentage variance explained by the top principal component of these 100 SWC time series was utilized as a test statistic (Zalesky et al., 2014). A high percentage variance would imply the existence of coherent SWC fluctuations across the 100 pairs of brain regions. The percentage variance computed from real data was compared against the null distribution from 1999 null datasets, resulting in one *p* value for each participant. Multiple comparisons across participants were corrected using a FDR of *q <* 0.05.

Figure 6A illustrates data from a representative HCP subject. Representative dynamic null data from PR, first order multivariate ARR and eleventh order bivariate ARR are shown in Figure 6B-D, while representative null data from a static null model (obtained by permuting the original fMRI time points) is shown in Figure 6E. The order of the bivariate ARR was chosen to match Zalesky et al. (2014). The top 100 most dynamic SWC time series (blue lines in Figure 6) exhibited massive fluctuations in the representative participant and corresponding null data. In the representative participant (Figure 6A), PR (Figure 6B), multivariate ARR (Figure 6C), bivariate ARR (Figure 6D) and static FC model (Figure 6E), the first principal component of the 100 SWC time series (red line in Figure 6) accounted for 62%, 58%, 66%, 11% and 19% of the variance respectively.

**Figure 6:**
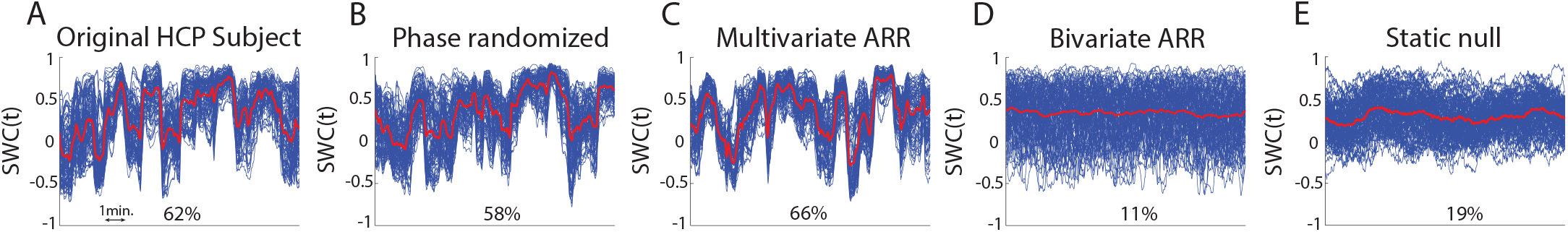
(A) Sliding window correlations (SWC) of a representative low motion HCP subject and example null data from (B) phase randomization (PR), (C) multivariate ARR, (D) bivariate ARR, and (E) static null model. The 100 most dynamic SWC time series are shown in blue, while their first principal component is shown in red. The percentage variance explained by the first principal component measures the coherence of brain dynamics. For the representative subject shown here, the percentage variance of the original data, PR, multivariate ARR, bivariate AR and static null model were 62%, 58%, 66%, 11% and 19% respectively. Therefore the coherence of brain dynamics was statistically indistinguishable among real data, PR and multivariate ARR, while bivariate ARR and static null data exhibited significantly less coherent brain dynamics.

On average (across 100 participants), the first principal component explained 49%, 46%, 45%, 10% and 27% variance in real data, PR, multivariate ARR, bivariate ARR and static null model respectively (Figure S2). For PR and multivariate ARR null data, the null hypothesis was rejected for only one participant. Therefore the stationary linear Gaussian model reproduces *coherent* SWC fluctuations for 99% of the low motion HCP participants. For bivariate ARR, the null hypothesis was rejected for all the participants. This discrepancy is discussed in the following section.

### 6.5. Why bivariate ARR might generate false positives

The previous two sections suggest that bivariate ARR commonly used in the literature (Chang and Glover, 2010; Zalesky et al., 2014) might be susceptible to false positives. To understand why this might occur, let us consider the toy example illustrated in Figure 7. In this toy example, there are three brain regions *X*, *Y* and *Z*, whose signals follow a first order AR model (Figure 7A). With the multivariate ARR procedure, the AR parameters were estimated using the time series from all three brain regions. The estimated AR parameters (Figure 7B) were the same as the true parameters (Figure 7A). As illustrated in Figure 7B, there is no arrow directly connecting brain regions *Y* and *Z*. In other words, brain regions *Y* and *Z* only influence each other via brain region *X*.

**Figure 7:**
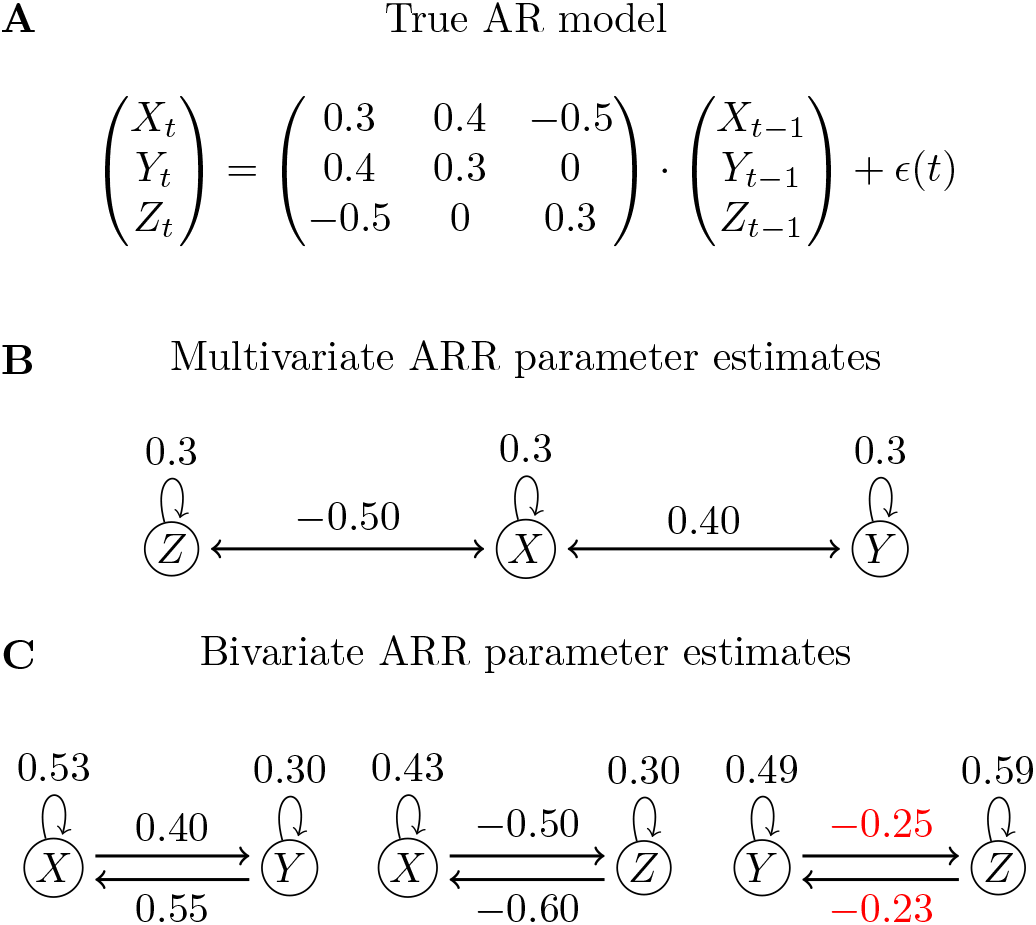
(A) Example AR model with three regions *X*, *Y* and *Z*. (B) AR model parameters estimated by considering the data from all three regions jointly (i.e., multivariate ARR). True AR parameters were recovered. (C) AR model parameters estimated for each pair of regions (i.e., bivariate ARR). Wrong AR parameters were recovered. In particular, direct interactions between regions *Y* and *Z* (red in figure) were found, which were non-existent in the true model.

On the other hand, the bivariate ARR procedure estimates AR parameters separately for brain regions *X* and *Y*, brain regions *X* and *Z*, and brain regions *Y* and *Z*. The estimated parameters (Figure 7C) are generally different from the true AR parameters (Figure 7A). More specifically, brain regions *Y* and *Z* exert direct inuence on each other (which is non-existent under the true model).

When generating null data using multivariate ARR, the time course at brain region *X* is generated by taking into account the inuence of both brain regions *Y* and *Z* (Figure 7B). However, in the bivariate ARR procedure, the time course at brain region *X* is generated by taking into account the inuence of only brain region *Y* (left panel of Figure 7C) or brain region *Z* (center panel of Figure 7C), but not both. Therefore bivariate ARR neglects inuence among all brain regions.

Furthermore, since bivariate ARR estimates the AR parameters for each pair of brain regions separately, any coherence among pairs of brain regions is destroyed. Consequently, the false positive situation appeared more severe when evaluating the existence of coherent brain dynamics (section 6.4) than when evaluating the existence of “dynamic” connections (section 6.3).

### 6.6. Control analyses

To ensure the results are robust to the particular choice of parameters, the sliding window size was varied from 20 frames to 100 frames, and the AR model order *p* was varied from 1 to 8. When evaluating coherence of whole brain dynamics, the number of most dynamic connections was also varied from 20 to 200. Since thresholding by DVARS might artificially exclude “real” dynamics, we also considered all the HCP subjects, rather than just the top 100 participants with least motion and DVARS. We also considered a higher resolution resting-state parcellation with 400 ROIs (Schaefer et al., 2017). None of these changes significantly affected the results.

Surprisingly, AR models of orders ranging from 1 to 8 explained similar variations in SWC fluctuations (Figure S3). However, AR model of order 0 (i.e., only preserving *R*_0_ or static FC) could not explain SWC fluctuations (Figure S3). Since our results might be sensitive to the choice of test statistic, the nonlinear statistic utilized in Zalesky et al. (2014) was also considered. We found that it was even more difficult to reject the null hypothesis using this nonlinear statistic.

The ICA-FIX fMRI data utilized in this work have been processed with a very weak highpass filter (2000s cut-off; Smith et al., 2013a). Additional highpass (0.0167Hz) filtering has been recommended to remove aliasing artifacts introduced by the SWC windowing procedure (Leonardi and Van De Ville, 2015). We found that the additional highpass filtering decreased the amplitude of the SWC fluctuations of both the original and surrogate data (Figure S4). Given that the windowing artifacts should also appear in the surrogate data, it is not obvious that applying highpass filtering would necessarily weaken statistical significance. However, in practice, highpass filtering did result in weaker statistical significance (Figure S4).

**Figure 8:**
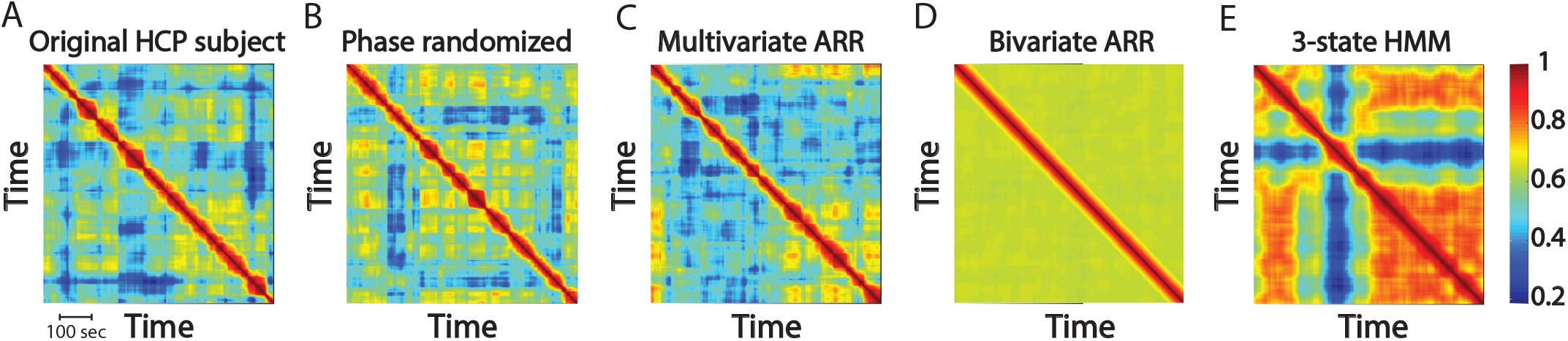
PR and multivariate ARR can replicate rich SWC dynamics of real data. Each plot is a *T* × *T* functional connectivity dynamics (FCD) matrix, where the *i*-th row and *j*-th column of the matrix corresponds to the correlation between the SWC of time points *i* and *j* matrix. (A) Representative HCP subject, (B) PR null data, and (C) first order multivariate ARR null data exhibit recurring SWC patterns. (D) Bivariate ARR null data exhibits weak recurring SWC patterns. (E) 3-state HMM null data exhibits sharp SWC transitions not present in original data. Here the FCD matrices were computed based on a parcellation into 114 ROIs (Yeo et al., 2011, 2015).

For completeness, we also applied bandpass filtering (0.01− 0.1Hz) to the data. Like highpass filtering, bandpass filtering decreased the amplitude of SWC fluctuations and weakened the statistical significance (Figure S4).

Finally, some authors have suggested that regressing mean grayordinate signal (akin to global signal regression) in addition to ICA-FIX might be necessary to remove global noise artifacts (Burgess et al., 2016; Siegel et al., 2016). When mean grayordinate signal was regressed, the SWC fluctuations were less statistically significant. For example, when evaluating the existence of coherent brain dynamics, the null hypothesis was not rejected for all 100 participants (Figure S4).

## 7. Stationary linear Gaussian models explain SWC fluctuations better than HMM

Section 6 suggests that distinguishing the linear stationary Gaussian model from real fMRI data is difficult (at least for the statistics tested). However, failure to reject the null hypothesis could be due to a lack of statistical power, rather than the null hypothesis being true. It is entirely possible that HMM-type models (Allen et al., 2014; Wang et al., 2016) might generate null data that fit observed fluctuations in SWC better than the linear stationary Gaussian model.

To test this possibility, Figure 8A shows the *T × T* functional connectivity dynamics (FCD) matrix of a representative HCP participant, where the *i*-th row and *j*-th column of the FCD matrix corresponds to the correlation between the SWC of time points *i* and *j* (Hansen et al., 2015). The presence of relatively large off-diagonal entries (yellow in Figure 8A) suggest the presence of recurring SWC patterns.

The PR (Figure 8B) and first order multivariate ARR (Figure 8C) null data were able to replicate the rich dynamics of the empirical FCD matrix (Figure 8A), while first order bivariate ARR null data (Figure 8D) exhibited significantly weaker recurring SWC patterns. By contrast, the 3-state HMM null data^11^ (Figure 8E) exhibited recurring SWC patterns with much sharper transitions than real data (Figure 8A).

To quantify these differences, the FCD matrices of the four null data generation approaches (Figure 8B-E) were compared with the empirical FCD matrix (Figure 8A) using the Kolmogorov-Smirnov statistic (Figure S5). Both the PR null data and (first order) multivariate ARR null data fitted the empirical FCD matrix better than the HMM null data (two-sample t-test *p <* 1*e*-32 and *p <* 1*e*-30 respectively).

11 states were necessary for the HMM model to perform as well as first order multivariate ARR (Figure S5). However, visual inspection of the FCD matrix (Figure S6A) suggests that the 11-state HMM still generated SWC patterns with sharper transitions than real data. For completeness, Figures S5 and S6B show the FCD results replicated with the Laumann null data generation approach (Laumann et al., 2016).

Finally, the results were replicated (Figure 9) in the one subject for whom the stationary, linear, Gaussian null hypothesis was rejected in Section 6.4. This illustrates the important point that rejection of the stationary, linear, Gaussian null hypothesis does not necessarily imply that the HMM would explain the fMRI data better than multivariate ARR or PR.

**Figure 9:**
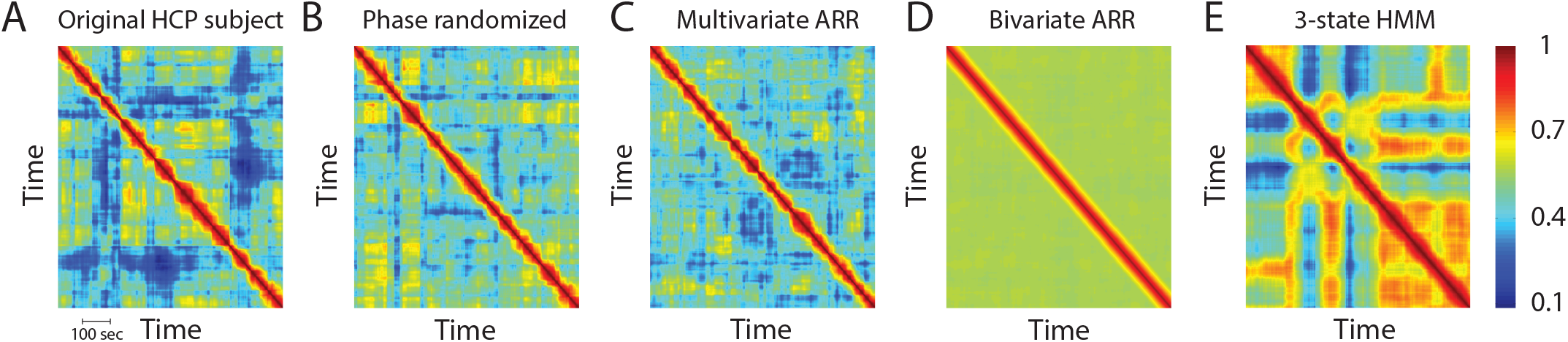
FCD matrices for (A) the one HCP subject for whom the stationary, linear, Gaussian null hypothesis was rejected in Section 6.4, (B) PR, (C) first order multivariate ARR, (D) bivariate ARR, and (E) 3-state HMM. Note the slight difference in scale as compared to Figures 8 and S6.

## 8. Discussion

In this article, we seek to improve our understanding of observed fluctuations in resting-state FC (e.g., SWC) widely reported in the literature. These fluctuations have often been interpreted as dynamic changes in inter-regional functional interactions, and non-stationary switching of discrete brain states (e.g., Allen et al., 2014). However, several recent papers have questioned these interpretations, especially in the case of single subject fMRI data (Hindriks et al., 2016; Laumann et al., 2016).

### 8.1. Linking Stationarity, dFC and brain states

By reviewing the conceptualization of fMRI as a random process (Section 3), we highlight that many statistical notions, such as *ensemble* auto-covariance and WSS, are reliant on ensemble statistics. Ensemble statistics are defined by taking into account an infinite number of realizations of a random process. However, the fMRI data of multiple participants are often considered as single realizations of different random processes. Therefore all FC measures in the literature are actually based on sample statistics (i.e., statistics based on one realization), but not ensemble statistics. It is possible that in the case of fMRI, ensemble statistics are equal to sample statistics. However, in this scenario, fMRI would be ergodic, which would in turn imply that fMRI is WSS.

Because the definition of WSS involves ensemble statistics, it is possible to come up with a toy brain with discrete brain states (i.e., HMM process) that is both WSS and ergodic (Section 4). Given that a WSS process can exhibit sharp transitions in SWC (Figure 2), this suggests that observed fluctuations in functional connectivity are not necessarily evidence of a non-stationary system.

The loose use of the term “non-stationarity” in the literature is not merely a linguistic issue, but can lead to potential confusion because current dFC statistical testing approaches rely on frameworks from the physics and statistics communities utilizing strict statistical notions, including stationarity (e.g., Schreiber and Schmitz, 2000). This motivates our detailed exploration of the assumptions behind the popular ARR and PR frameworks for generating null data for hypothesis testing of dFC.

It is possible that the dFC community might not be referring to “non-stationarity” in the statistical sense. However, the widely used null hypothesis testing frameworks (ARR and PR) do rely on traditional statistical notions of stationarity. Therefore it is important for the community to articulate the exact statistical notions (e.g., piecewise stationarity; Nason (2013)) that might encode the intuitive notion of “non-stationarity” mentioned in the dFC literature. Having the exact statistical notions will lead to better null hypothesis testing frameworks.

### 8.2. Preserving auto-covariance beyond static FC

Our review of ARR and PR frameworks (Section 5) shows that both approaches retain the 0-th sample auto-covariance (*R*_0_ in Eq (2)) of the original fMRI data. *R*_0_ can be interpreted as *static* FC (being an unnormalized variant of Pearson’s correlation). Therefore *R*_0_ is an important quantity to preserve in null data since the dFC researcher is presumably interested in showing that dFC cannot be completely explained by static FC.

Preserving *R*_0_ (static FC) can be easily achieved by permuting the temporal ordering of fMRI time points. However, such a procedure ignores well-known auto-correlation in the original fMRI data, which will lead to false positives in null hypothesis testing. Both PR and ARR seek to preserve auto-correlations in addition to *R*_0_. More specifically, PR preserves the entire *sample* auto-covariance sequence (*R*_0_,*… R_T −_*_1_ in Eq (2)) of the original fMRI data, while ARR of order *p* preserves only the first *p* + 1 terms of the auto-covariance sequence (*R*_0_, *… R_p_*). For example, ARR of order 1 preserves *R*_0_ and *R*_1_.

Since PR potentially preserves higher order auto-covariances than ARR, it might be theoretically advantageous. However, in our experiments, PR and multivariate ARR of order 1 were able to explain observed SWC in real fMRI data equally well (Figures 5, 6 and 8). On the other hand, ARR of order 0 (i.e., only *R*_0_ or static FC is preserved) does not explain SWC fluctuations at all (Figure S3). The implication is that fluctuations in SWC can largely be explained by taking into account auto-covariances of lag 0 (i.e., static FC or *R*_0_) and lag 1 (i.e., *R*_1_) from the original fMRI data.

### 8.3. Null data generation in the literature

Understanding why ARR and PR are able to preserve sample auto-covariances of the original data (Figure 3) is useful for interpreting other null data generation approaches in the literature. For example, Allen et al. (2014) applies the PR framework directly on the sliding window FC time series, and not on the fMRI time series. Therefore in the case where the same random phase sequence was added to all dFC phase spectra (i.e., SR1 in Allen et al., 2014), the null dFC time series have the same auto-covariance sequence as the original dFC time courses, but not the auto-covariance of the original fMRI time courses. Therefore the tested null hypothesis is similar, but not the same as other papers that applied PR to the fMRI time series (Handwerker et al., 2012; Hindriks et al., 2016).

ARR has also been widely used in neuroimaging applications, mostly following the bivariate variant proposed by Chang and Glover (2010). Our results suggest that bivariate ARR neglects higher-order interactions (Figure 7), resulting in wrongly estimated AR model parameters, potentially leading to false positives. The issue with wrongly estimated AR model parameters is reminiscent of Friston’s criticism of functional connectivity (Friston, 2011), where two regions A and B might be functionally connected because of mutual effective connectivity with an intermediary region C. However, it should be noted that AR models are not effective connectivity models because of the lack of hemodynamic modeling (Friston, 2009). The false positive rate associated with bivariate ARR is less serious when investigating the existence of individual “dynamic” edges (Figure 5), but extreme when investigating the coherence of SWC across dynamic pairs of brain regions. More specifically, when investigating SWC coherence, the null hypotheses for all participants were rejected using bivariate ARR, compared with only one participant for multivariate ARR or PR (Figures 6 and S2).

More recently, Laumann et al. (2016) proposed a procedure to generate null data that matches the static FC (*R*_0_) and the power spectral density of the original fMRI data (averaged across ROIs). Given the deep relationship between the auto-covariance sequence and cross-spectral density of the original fMRI data (Appendix A5), preserving the average power spectral density retains some (but not all) information of the original fMRI auto-covariance structure *beyond* static FC. The advantage of the Laumann (and PR) approaches over multivariate ARR is that the number of fitted parameters does not increase quadratically with the number of ROIs. This allows the generation of null data with large number of ROIs for which multi-variate AR model parameters cannot be estimated (see Appendix A2). However, because the Laumann approach preserved fewer properties of the original data than PR or multivariate ARR, one might expect the Laumann null data to be less close to real data than PR or multivariate ARR. Indeed, this expectation is empirically confirmed in Figures S5 and S6B. On the other hand, these results also suggest that compared with bivariate ARR, the Laumann procedure generated null data that were more similar to real data. This explains why Laumann et al. (2016) has more difficulties rejecting the null hypothesis compared with Chang and Glover (2010) and Zalesky et al. (2014).

Another approach of generating null data exploits the approximate scale-free or scale-invariance nature of fMRI. More specifically, the fMRI power spectrum is known to approximately follow a power law distribution over the frequency band spanning 0.0005Hz to 0.1Hz (Ciuciu et al., 2012; He, 2011, 2014). The upper limit of this frequency band reflects the lowpass characteristics of neurovascular coupling (Hathout et al., 1999; Anderson, 2008), while the lower limit reflects the maximum practically achievable duration of continuously awake scanning, which is about 30 minutes.

The exponent of the power law distribution can be used to inform fMRI stationarity (Eke et al., 2002; He, 2011; Ciuciu et al., 2012). Hypothesis testing can be performed by comparing real data to null, stationary, scale-free data with matched power law exponent. Failure to reject the null hypothesis indicates scale-invariance and stationarity. Across a variety of sophisticated approaches, including wavelet leaders, resting-state fMRI has been found to be stationary (He, 2011; Ciuciu et al., 2012), whereas electrical field potentials and electroencephalograms have been found to be non-stationary (Miller et al., 2009; Milstein et al., 2009; Freeman and Zhai, 2009; He et al., 2010; Van de Ville et al., 2010). One key difference between the scale-free literature and the PR/ARR framework explored in this article is that the former has mostly focused on a single time series from an individual ROI (or voxel or functional network), while the latter models multivariate interactions between ROIs.

As we focus in this paper on the importance of considering the auto-covariance structure of fMRI time series (Eq. (2)), the Wiener-Khintchine theorem (Appendix A5) provides a link to the scale-free literature by detailing the relationship between the power spectrum and auto-correlation. Based on this, one can for example interpret a larger power-law exponent as an indicator of stronger auto-correlation, and vice versa. Intriguingly, the power law exponent is smaller during task engagement and larger during drowsiness and sleep (He and Raichle, 2009; He, 2011; Ciuciu et al., 2012, 2014; Churchill et al., 2016).

### 8.4. The stationary, linear, Gaussian null hypothesis

Our review suggests that PR and ARR generate null data that are linear, WSS and Gaussian (Section 5). Conversely, any linear Gaussian process can be arbitrarily well approximated by an AR model of sufficiently large order *p*. Together, this implies that if the original time series significantly differ from ARR or PR null data, then the original data is non-Gaussian or nonlinear or non-stationary. For example, the null hypothesis was easily rejected for the stationary two-state toy brain (Figure 4) because the toy brain is nonlinear and non-Gaussian.

Nevertheless, this ambiguity is less of an issue because our results also suggest that the stationary linear Gaussian null hypothesis cannot be rejected for most low-motion HCP participants (Section 6). On average (across participants), the stationary linear Gaussian null hypothesis can only be rejected for 0.6% of brain region pairs (Figure 5) when using PR and multivariate ARR. When studying dFC coherence, the stationary linear Gaussian null hypothesis was only rejected for 1% of the participants (Figures 6, S2 and S3).

The difficulty in rejecting the null hypothesis is somewhat surprising given that the brain is a complex organ possessing nonlinear neuronal dynamics (e.g., Hodgkin and Huxley, 1952; Valdes et al., 1999; Deco et al., 2008; Stephan et al., 2008; Deco et al., 2011). However, our results are consistent with previous literature reporting difficulties to reject the null model, especially in single subject fMRI data (Hindriks et al., 2016; Laumann et al., 2016). A close look at the dFC literature suggests similar difficulties in seminal dFC papers. For example, dFC states were found using null data generated by adding the same random phase sequence to all dFC phase spectra during PR (i.e., SR1 in Allen et al., 2014). Similarly, Zalesky and colleagues (Zalesky et al., 2014) reported that on average across subjects, the null hypothesis was rejected for only 4% of edges when using bivariate ARR, which is highly consistent with our bivariate ARR results, where 5% of edges were rejected. Therefore, the dFC literature is consistent in finding only small deviations from the stationary, linear, Gaussian null hypothesis. Much of the controversy might be due to differences in the interpretation, i.e., viewing the glass as half-full or half-empty.

For the small minority of edges or participants whose null hypothesis is rejected, it is unclear whether this deviation is due to non-stationarity, nonlinearity or non-Gaussianity. It is also unclear whether these deviations are due to artifacts (e.g., respiration; Laumann et al., 2016; Power et al., 2017) or neurologically meaningful. One approach of demonstrating neurological relevance is by association with behavior or disease. While there are many studies linking dFC measures (e.g., SWC, dwell time of brain states, etc) with behavior and diseases (Damaraju et al., 2014; Barttfeld et al., 2015; Su et al., 2016; Du et al., 2016; Nomi et al., 2017; Shine et al., 2016; Wang et al., 2016), there are far fewer studies explicitly demonstrating that dFC measures are able to explain behavioral measures or disease status above and beyond static FC (e.g., Rashid et al., 2014). Moreover, to the best of our knowledge, we are unaware of any studies showing dFC-behavioral associations above and beyond the stationary, linear and Gaussian model. Therefore, for studies demonstrating that their dFC measures are more strongly associated with behavior than static FC (e.g., Rashid et al., 2014), the improvement might potentially be explained by the AR model, which encodes both static FC (*R*_0_) and linear dynamical interactions between brain regions (*R*_1_, etc).

### 8.5. Does dynamic functional connectivity exist?

Does stationarity, linearity and Gaussianity of fMRI time series imply that dFC is spurious? Obviously, if dFC is strictly defined as non-WSS (Section 3.2), then stationarity does imply the lack of dFC. However, Section 4 suggests that such a definition of dFC would exclude a class of signals (e.g., HMM) that most neuroscientists would think of as encoding dFC. Therefore alternative definitions of dFC should be considered.

If dFC is thought of as corresponding to the brain sharply switching between discrete states with distinct FC patterns, then our results suggest a lack of evidence in resting-state fMRI. The presence of HMM-type states could potentially lead to the rejection of the stationary linear Gaussian null hypothesis (Section 4). However, the null hypothesis was not rejected for most low motion HCP participants (Section 6). This non-rejection could be due to a lack of statistical power. Therefore we tested whether an HMM-type model explicitly encoding the presence of states would fit SWC fluctuations better than AR (or PR) models. We found that ARR and PR reproduced the gentle fluctuations of recurring SWC patterns in real data, whereas sharp transitions were observed in the HMM (Figure 8). The result was replicated (Figure 9) even in the one subject for whom the stationary linear Gaussian model was rejected^12^ (Section 6.4). Altogether, this suggests the lack of discrete brain states as measured by SWC, consistent with some proposals that SWC might be better explained by a mixture of states (Leonardi et al., 2014; Miller et al., 2016). One difficulty in estimating discrete FC states with SWC is that SWC introduces additional blurring of the fMRI signal, which is already a smoothed response to underlying neural signal. However, it is worth noting that a recent approach applying hemodynamic de-convolution and clustering also yielded overlapping activity-level states without sharp temporal switching (Karahanoğlu and Van De Ville, 2015).

If dFC is thought of as the existence of FC information beyond static FC, then our results do support the existence of dFC because multivariate AR models explained SWC fluctuations significantly better than just static FC (Figure S3). Indeed, AR models are often considered models of linear dynamical systems (e.g., Casti, 1986; Gajic, 2003). By encoding linear dynamical interactions between brain regions (*A*_1_ in Eq. (3)), the first order AR model captures both static FC (i.e., *R*_0_ in Eq. (2)) and dynamic FC structure (i.e., *R*_1_ in Eq. (2)). It is worth reminding the readers that the diagonal elements of *R*_1_ encode the auto-correlation within individual brain regions, while the off-diagonal terms encode lagged cross-covariance between brain regions. Since the Laumann null data generation approach (Laumann et al., 2016) explicitly preserves static FC and temporal auto-correlation, but did not explain SWC as well as first order AR models (Figure S6), this suggests the importance of the off-diagonal entries of *R*_1_, i.e., lagged cross-covariance between brain regions. The importance of such resting-state lagged cross-covariances has been described in humans (Mitra et al., 2014; Raatikainen et al., 2017) using fMRI and in animals using a variety of electrophysiological techniques (Mohajerani et al., 2013; Stroh et al., 2013; Matsui et al., 2016); for review, see Mitra and Raichle (2016).

Finally, if dFC refers to the presence of biological information in FC fluctuations, then recent evidence suggests that FC fluctuations within a single fMRI session can be linked to varying levels of vigilance (Wang et al., 2016). Others have shown that activity level fluctuations might also be associated with arousal (Chang et al., 2016), attention (Kucyi et al., 2016) or emotional states (Kragel et al., 2016). However, it is currently unclear whether the mental fluctuations might be more readily explained by fluctuating activity level (first order statistics) or fluctuating FC (second order statistics).

It is important to emphasize that stationary, linear, Gaussian fMRI does not contradict the presence of mental fluctuations during resting-state fMRI. To see this, let us consider the following toy example. Suppose Alice and Bob play a game, where Alice tosses a fair coin at each round of the game. Every time the coin toss results in a head, Alice pays Bob a dollar. Every time the coin toss results in a tail, Bob pays Alice a dollar. We can see that the coin tosses do not possess any interesting dynamics: coin tossing is stationary and temporally independent (so it is even less “interesting” than fMRI). However, the outcomes of the coin toss are still financially (behaviorally) relevant. Similarly, stationary, linear, Gaussian fMRI dynamics might reflect fluctuation in the levels of vigilance, arousal, attention and/or emotion (Chang et al., 2016; Kucyi et al., 2016; Kragel et al., 2016; Wang et al., 2016).

### 8.6. Future directions

Rather than arguing about the existence of dFC, it might be more useful to re-frame this debate in terms of adequate models of fMRI time series: (i) what kind of model reproduces properties of fMRI (and dFC) time series time series and (ii) what types of FC fluctuations (shapes, timescales) are expected in these models? This framing sidesteps the question of whether `dFC exists’, but instead relies on mathematical models, from which principled predictions and interpretations might provide further insights into human brain organization.

Given the ability of AR (and PR) models to generate realistic SWC, rather than treating them as just null models, AR models could themselves be utilized to provide insights into the brain. Since more complex models are harder to interpret, they should not be preferred unless existing models could not fit some important aspect of fMRI data. For example, since models of static FC cannot explain SWC dynamics very well (Figure S5), it is clear that first order multivariate AR model is preferable for a researcher interested in dFC. The next step would be to show that the additional dynamics modeled by AR model (above and beyond static FC) is functionally meaningful, such as by association with behavior or disease.

Since this article only focuses on SWC and several statistics (Sections 6.3, 6.4 and 7), it is possible that stationary linear Gaussian (SLG) models are unable to explain unexamined aspects of fMRI dynamics captured by non-SWC approaches or other statistics (Maiwald et al., 2008; Griffa et al., 2017). In these cases, more complex generative models, such as those involving mixture of states (Leonardi et al., 2014) or wavelets (Breakspear et al., 2004; Van De Ville et al., 2004), might be necessary. Generalizations of the AR model allowing for time-varying properties of their parameters, such as the GARCH model (Bollerslev, 1986) might also be considered (Lindquist et al., 2014; Choe et al., 2017). Additional examples of more advanced approaches can be found in the excellent review by Preti and colleagues (Preti et al., 2016).

However, it is worth emphasizing that following Occam’s razor, more complex models should be used only if simpler ones have been proven to fail at capturing important information encoded in the data. In this paper we have shown that the stationary linear Gaussian model captures information beyond classical static models of FC, but it is not clear whether more complex and more biophysically realistic models will further explain fMRI dynamics. For example, the use of the FCD matrices (Figure 8) to visualize the rich SWC dynamics was pioneered by Hansen et al. (2015), who demonstrated that nonlinear biophysical (neural mass) models could replicate some of the rich SWC dynamics, while a linear dynamical model was not able to. The linear dynamical model utilized by Hansen and colleagues is essentially the same as the first order AR model utilized here. However, the linear dynamical model was utilized to model neural dynamics, rather than fMRI data directly. The “output” of the linear neural model was then fed into a biophysical hemodynamic response model to generate fMRI data. Importantly, the interactions between brain regions were set to be the diffusion connectivity matrix, rather than fitted to real fMRI data. Conse-ries and (ii) what types of FC fluctuations (shapes, timescales) quently, the linear dynamical model (Hansen et al., 2015) are expected in these models? This framing sidesteps the question of whether ‘dFC exists’, but instead relies on mathematical models, from which principled predictions and interpretations might provide further insights into human brain organization. did not exhibit the rich dynamics observed in our experiments (Figure 8). Because of the difficulties in estimating the parameters of the nonlinear neural mass models, it is unclear whether an optimized neural mass model might be able to explain FC fluctuations better than a first order multivariate AR model.

A recurring criticism of AR modeling of fMRI time series is that the model parameters cannot be interpreted as effective connectivity because spatial variability of the hemodynamic response function is usually not taken into account (e.g., Friston, 2009). Hence other generative models such as dynamic causal modelling should be preferred if one wants to study effective connectivity (Friston et al., 2003). However, this does not preclude the use of AR models as a diagnostic tool encoding functional connectivity information beyond static FC (Rogers et al., 2010). An obvious challenge will then be to extract the most relevant information from these models (Liégeois et al., 2015; Ting et al., 2016).

## 9. Conclusion

We explore statistical notions relevant to the study of dynamic functional connectivity. We demonstrate the existence of a stationary process exhibiting discrete states, suggesting that stationarity does not imply the absence of brain states. Our review of two popular null data generation frameworks (PR and ARR) suggests that rejection of the null hypothesis indicates non-stationarity, nonlinearity and/or non-Gaussianity. We show that most HCP participants possess stationary, linear and Gaussian fMRI during the resting-state. Furthermore, AR models explain real fMRI data better than just static FC, and a popular approach that explicitly models brain states. Overall, the results suggest a lack of evidence for discrete brain states (as measured by fMRI SWC), as well as the existence of FC information beyond static FC. Therefore dFC is not necessarily spurious because AR models are themselves linear dynamical models, encoding temporal auto-covariance above and beyond static FC. Given the ability of AR models to generate realistic fMRI data, AR models might be well suited for exploring the dynamical properties of fMRI. Finally, our results do not contradict recent studies showing that temporal fluctuations in functional connectivity or activity level can be behaviorally meaningful. The code used for this work are available at GITHUB_LINK_TO_BE_ADDED.

## Acknowledgements

The authors would like to thank Andrew Zalesky, Tom Nichols and the anonymous reviewers for their insightful feedback. The authors would also like to thank the participants of the interesting twitter threads on this topic (https://twitter.com/russpoldrack/status/879360707095023616 and https://twitter.com/danjlurie/status/891113274586025984), which have helped to shape certain subtle points in this work. This research was supported by Singapore MOE Tier 2 (MOE2014-T2-2-016), NUS Strategic Research (DPRT/944/09/14), NUS SOM Aspiration Fund (R185000271720), Singapore NMRC (CBRG/0088/2015), NUS YIA, Singapore NRF fellowship (NRF-NRFF2017-06), the Neuroimaging Informatics and Analysis Center (1P30NS098577), F30MH100872 (T.O.L.), and by a W.B.I.- World Excellence Fellowship (R.L.). Data were provided by the Human Connectome Project, WU-Minn Consortium (Principal Investigators: David Van Essen and Kamil Ugurbil; 1U54MH091657) funded by the 16 NIH Institutes and Centers that support the NIH Blueprint for Neuroscience Research; and by the McDonnell Center for Systems Neuroscience at Washington University.

## Appendix

### A1. Details of ARR procedure and linear Gaussianity

Let *x_t_* be the *N* × 1 vector of fMRI data at time *t* after each time course has been demeaned. There are many approaches to estimate the *p*-th order AR model parameters (Σ, *A*_1_, *A_p_*) from the data *x_t_*. A common approach is as follows.

1. Estimate *A*_1_,*…, A_p_* using the following least-squares cost function:

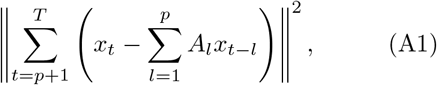

where *T* is the number of time points. Let *A* = [*A*_1_,*… A_p_*] (i.e., *A* is an *N* × *N p* matrix). Then the optimum to the above criterion corresponds to

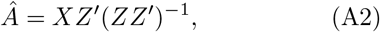 Where ’indicates transpose, *X* is an *N ×* (*T − p*) matrix

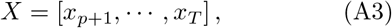

and *Z* is an *N p ×* (*T − p*) matrix

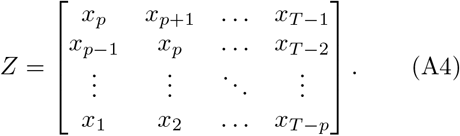 The estimation procedure therefore requires *ZZ^’^* (in Eq. (A2)) to be full rank, which implies that *T ≥* (*N* + 1)*p*. For example, if one utilized a parcellation with 400 ROIs, then one would need at least 401 *×* 3 = 1203 time points for a 3rd order AR model.
2. After estimating *Â*= [*Â_1_*,…, *Â_p_*], the residual (error) is defined as follows

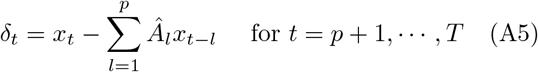

The covariance matrix Σ can then be estimated via the empirical covariance matrix:

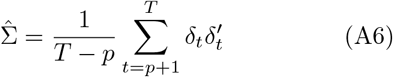

Once (Σ, *A* _1_, …, *A_p_*) are estimated, each ARR null data is initialized by randomly selecting *p* consecutive time points of the original data, and then repeatedly applying the AR model (Eq. (3)) until null data of length *T* are generated. By virtue of the AR model (Eq. (3)), the resulting null data is linear Gaussian if the resulting system is stable (see Appendix A2).

In the fMRI literature, it is common to generate null data where *∈*_t_ in Eq. (3)) are not generated using a Gaussian distribution, but obtained by sampling from the residuals *δ_t_* of the identified AR model (Chang and Glover, 2010). In practice, the residuals might not follow a Gaussian distribution (Lutkepohl, 2005), and so the resulting ARR null data would only be linear but not Gaussian. However, in our experiments (not shown), this difference does not have any practical effect on our conclusions.

### A2. WSS, stability and auto-covariance of ARR null data

The AR random process (Eq. (3)) is WSS if and only if the AR model is stable (Lutkepohl, 2005; Zivot and Wang, 2006; Pfaff, 2008). Stability can be assessed by ensuring that the matrix

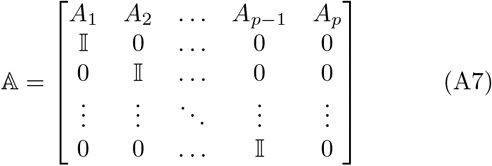

has eigenvalues with magnitude strictly smaller than one.

Therefore ARR null data is WSS assuming that the estimated AR parameters correspond to that of a stable AR model. This stability condition should therefore be checked when estimating the AR model parameters (Eq. (A1)). Since the original fMRI data is stable (i.e., the fMRI measurements do not diverge to infinity), the estimated AR model should be stable as long as the estimation procedure is reliable. In our experience with fMRI data, this is indeed the case if the AR order *p* is not too close to its maximal value of 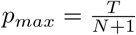 (Appendix A1). If the order *p* is too close to *pmax*, then the estimation procedure (Eq. (A1)) might require the inversion of an almost singular matrix, which might lead to an unstable AR model.

We now turn our attention to the relationship between the AR model parameters (Σ, *A*_1_, *…, A_p_*) (Eq. (3)) and auto-covariance sequence *R*_0_, *R_p_* (Eq. (2)), which is governed by the Yule-Walker equations (Yule, 1927; Walker, 1931) assuming infinitely long time courses:

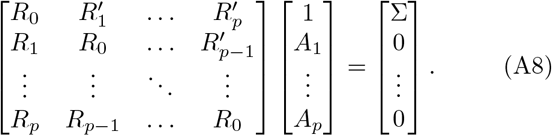

Eq. (A8) is invertible for sufficiently large *T*, and so the AR model parameters are completely determined by the auto-covariance sequence of the original data (Stoica and Moses, 2005) and vice versa (e.g., Pollock, 2011, Chapter 13). Consequently, the AR null data generated by this framework share the first *p* + 1 auto-covariances (Eq. (2)) computed from the original data.

It is also worth noting that for sufficiently large *T*, the AR parameters obtained by inverting Eq. (A8) are mathematically and practically equivalent to those obtained from Appendix A1. For small *T*, AR model parameters estimated from Eq. (A8) are guaranteed to correspond to a stable system, while AR model parameters estimated from Eq. (A2) are not (Stoica and Nehorai, 1987). Therefore one should always check the stability of the AR system when using the least squares approach (Appendix A1).

### A3. Details of the PR procedure

This appendix elaborates the PR procedure. Let *x^n^* denote the time course of the *n*-th brain region, while 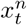 denote the *t*-th time point of the *n*-th brain region (where the first time point corresponds to *t* = 0). Without loss of generality, we assume that the time courses have been demeaned. PR proceeds as follows:

1. The Discrete Fourier Transform (DFT) for each time course *x^n^* is computed:

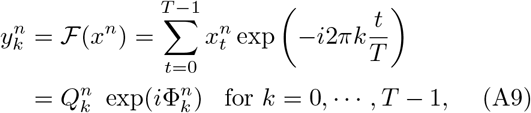

where *k* indexes frequency, and 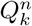 and 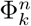 are the amplitude and phase of the *k*-th frequency component of the DFT. Since the input signal is real, 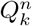 = 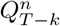 and 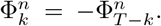 Because the signal has been demeaned, therefore 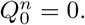.
2. A random phase *φ_k_* is then added to the DFT coefficients for each brain region *n*:

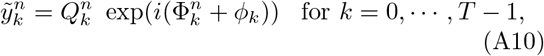

where *φ_k_* is drawn from a uniform distribution on the [0, 2*π*] interval. Importantly, *φ_k_* is the same for all brain regions and independently sampled for frequencies *k* = 0, *· · ·, 1T /*2*1* (although we note that *φ*_0_ is useless because *\*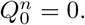*). For *k >[T/2]* (*[·]*denote the ceiling function), *φ_k_* = *−φ_T −k_* because of the need to ensure the null data remains real (rather than complex-valued).
3. The inverse DFT is then performed for each brain region *n* resulting in the PR null data:

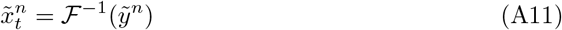

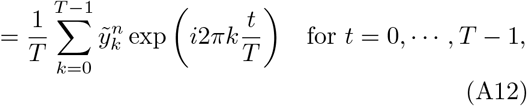 Because 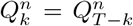, 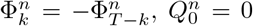, and *φ_k_* = *φ_T −k_* for *k > [T /*2], the null data (Eq. (A12)) can be written in the following form:

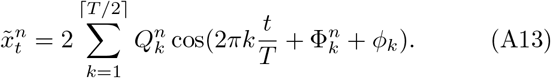

### A4. PR null data are WSS

To show that the PR null data is WSS, we show that the ensemble mean and ensemble auto-covariance do not depend on the time *t*.

First, by applying “expectation” to Eq. (A13) and via the linearity of expectation, the ensemble mean is equal to:

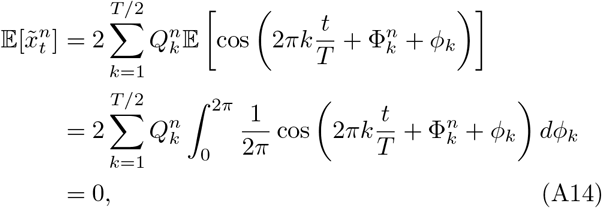

where the second equality arises because *φ_k_* ~ U[0, 2*π*]. Therefore the ensemble mean does not depend on the time *t*.

Let 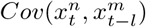 be the ensemble auto-covariance between the signal of brain region *n* at time *t* and the signal of brain region *m* at time *t–l*. Because the ensemble mean is equal to 0 for all time points, therefore

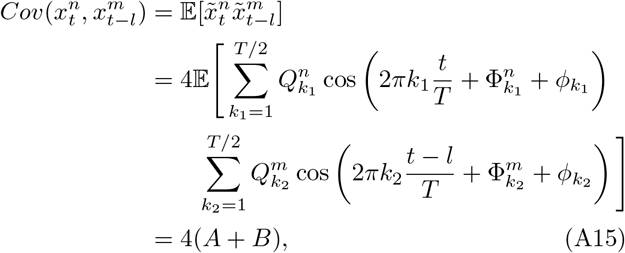

where *A* corresponds to the products of cosines with different frequencies and *B* corresponds to the products of cosines with the same frequencies. More specifically,

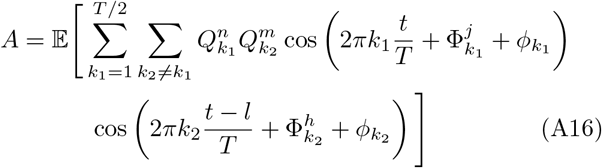

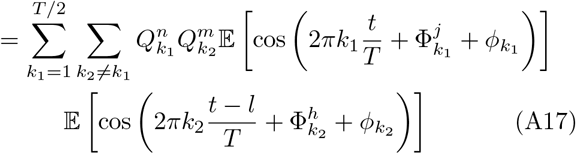

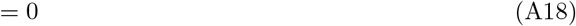

where the second equality is true due to the fact that the random phases *φ_k_*_1_ and *φ_k_*_2_ are independently sampled, and the third equality is obtained from straightforward integration (just like the ensemble mean). On the other hand,

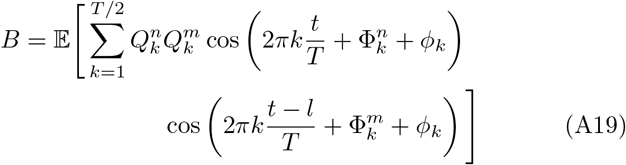

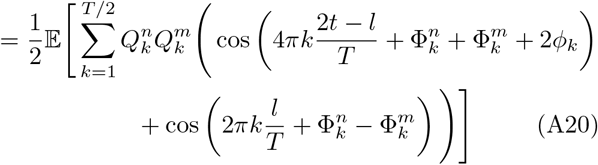

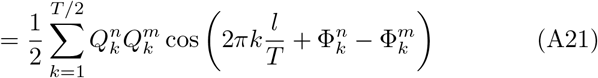

Therefore

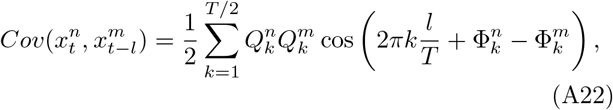

which does not depend on time *t* and only depends on the interval *l* between the two time points *t* and *t–l*. Therefore PR null data are WSS. Furthermore, one can also verify that the sample mean is equal to 0 and the sample auto-covariance sequence *R_l_* are equal to the ensemble auto-covariances (Eq. (A22)). Therefore PR null data are also ergodic.

### A5. PR null data preserve auto-covariance sequences

In this appendix, we explain why PR null data preserve auto-covariance sequences (Eq. (2)) of the original data. This property arises from the Wiener-Khintchine theorem, first formulated in the univariate case by Wiener (1930) and Khintchine (1934), and then reported in the multivariate case as an extension of the Wiener-Khintchine theorem (Prichard and Theiler, 1994), or cross-correlation theorem (Weisstein, 2016).

Given two time courses 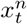 and 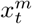 from brain regions *n* and *m*, their (sample) cross spectral density (CSD) is defined as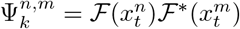, where *F*(·) is the DFT, *k* indexes frequency, and *∗* is the complex conjugate. Let 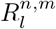 correspond to the *n*-th row and *m*-column of the auto-covariance matrix *R_l_* defined in Eq. (2). Then according to the multivariate Wiener-Khintchine theorem:

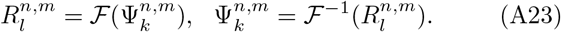

In other words, the (sample) CSD and the auto-covariance sequence of a multivariate random process encode the same information about the data.

Let the PR null time courses for brain regions *n* and *m* be denoted as 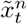 and 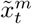 Their sample CSD corresponds to

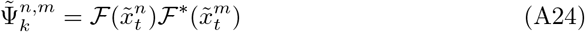

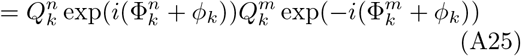

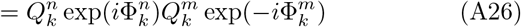

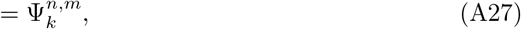

where the second equality is obtained by plugging in Equation (A10), and the random phase *φ_k_* is the same for all brain regions and therefore cancels out in the third equality. Since the sample CSD is the same between brain regions *n* and *m* in the PR null data, then their auto-covariance sequence is also the same according to Wiener-Khintchine theorem (Eq. (A23)).

## Supplementary material

### s1. Rejection of the stationary linear Gaussian model does not imply presence of states

Figure S1 (top) shows a representative time course generated from the following univariate model:

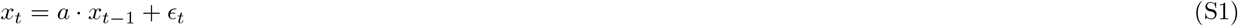

where *a* = 0.2 and *∈_t_* is noise with uniform distribution: *∈_t_* ~ *U* [−0.5, 0.5]. This process is stationary, linear and *non*-Gaussian. It is also clear that there is no discrete state associated with this process.

**Figure S1:**
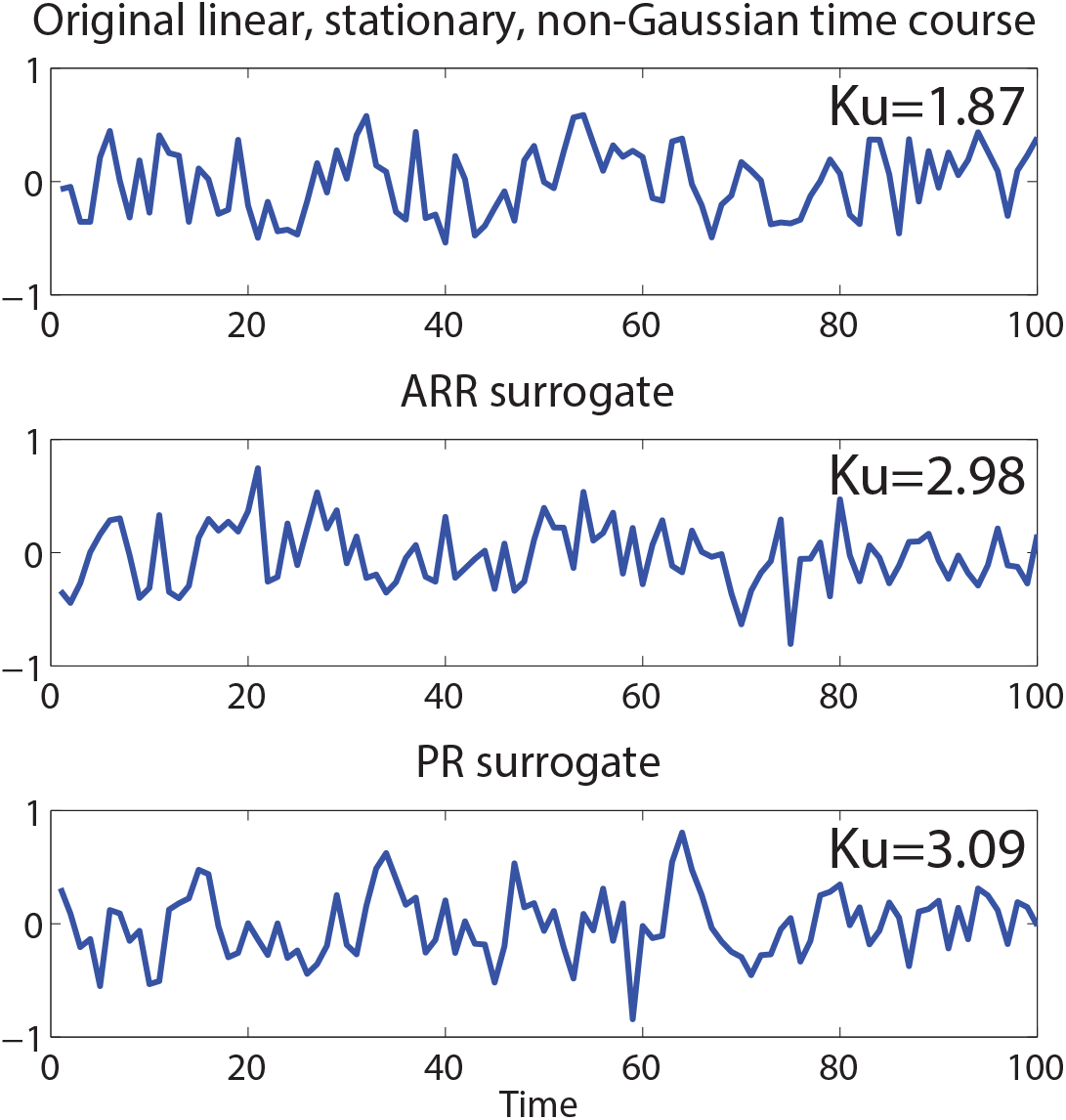
The stationary linear Gaussian model can be rejected because of non-Gaussianity of the original time course. (A) Original time course (top), one ARR surrogate (middle) and one PR surrogate with kurtosis (bottom). Kurtosis (*Ku*) of the original time course is 1.87 and Kurtosis of the sample surrogates are 2.98 (ARR) and 3.09 (PR).

PR and ARR were utilized to generate 999 surrogates (Figure S1). Kurtosis (*Ku*) was utilized as a test statistic (Figure S1). The kurtosis of the original time course is 1.87. The kurtosis of the surrogate time courses ranges from 2.43 to 4.10 (for ARR) and 2.31 to 4.36 (for PR). The stationary, linear, Gaussian null model was rejected with *p <* 0.001.

### S2. Coherence of SWC dynamics

**Figure S2:**
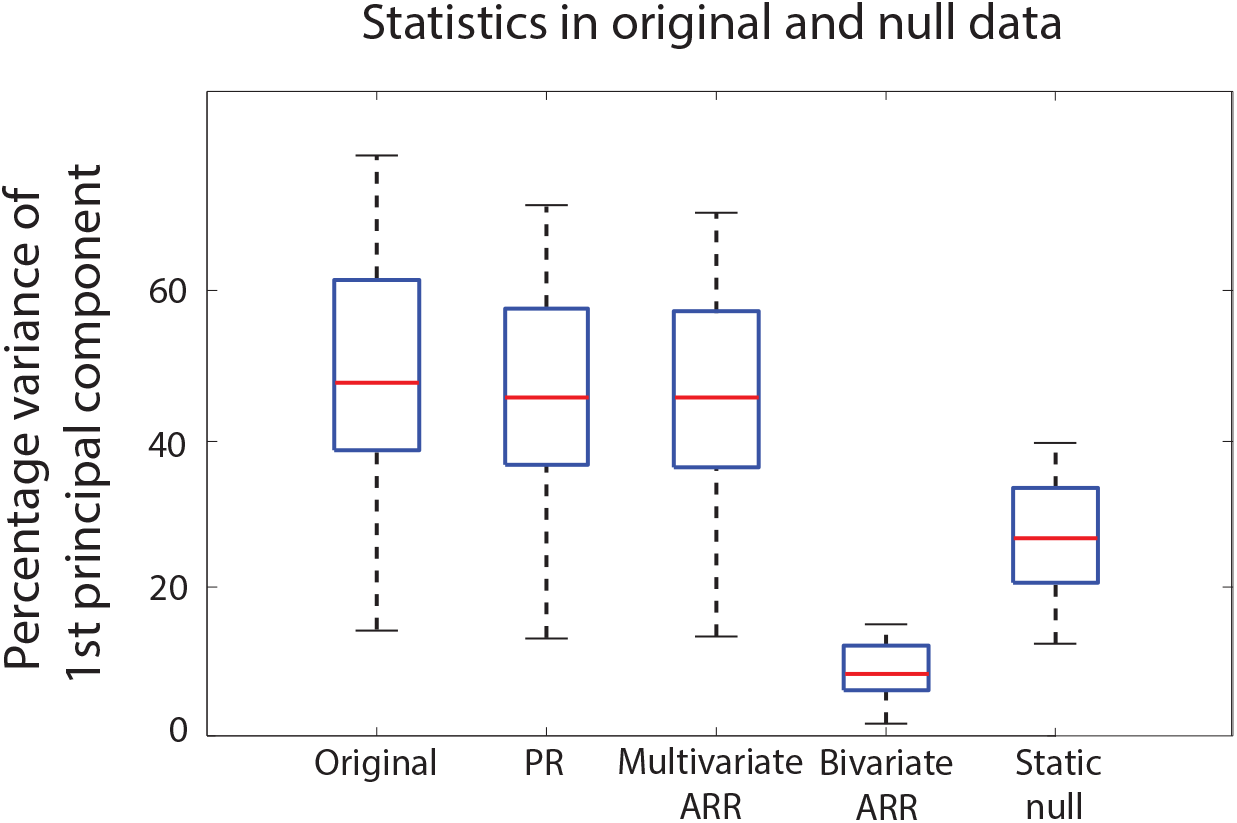
Distribution of percentage variance of first principal component across 100 participants in original, phase randomized (PR), multivariate ARR, bivariate ARR, and static null data. For 99% of the subjects, the stationary linear Gaussian null-model cannot be rejected using a false discovery rate of *q <* 0.05. (12 subjects had an uncorrected p value of less than 0.05).

### S3. Impact of ARR order *p*

**Figure S3:**
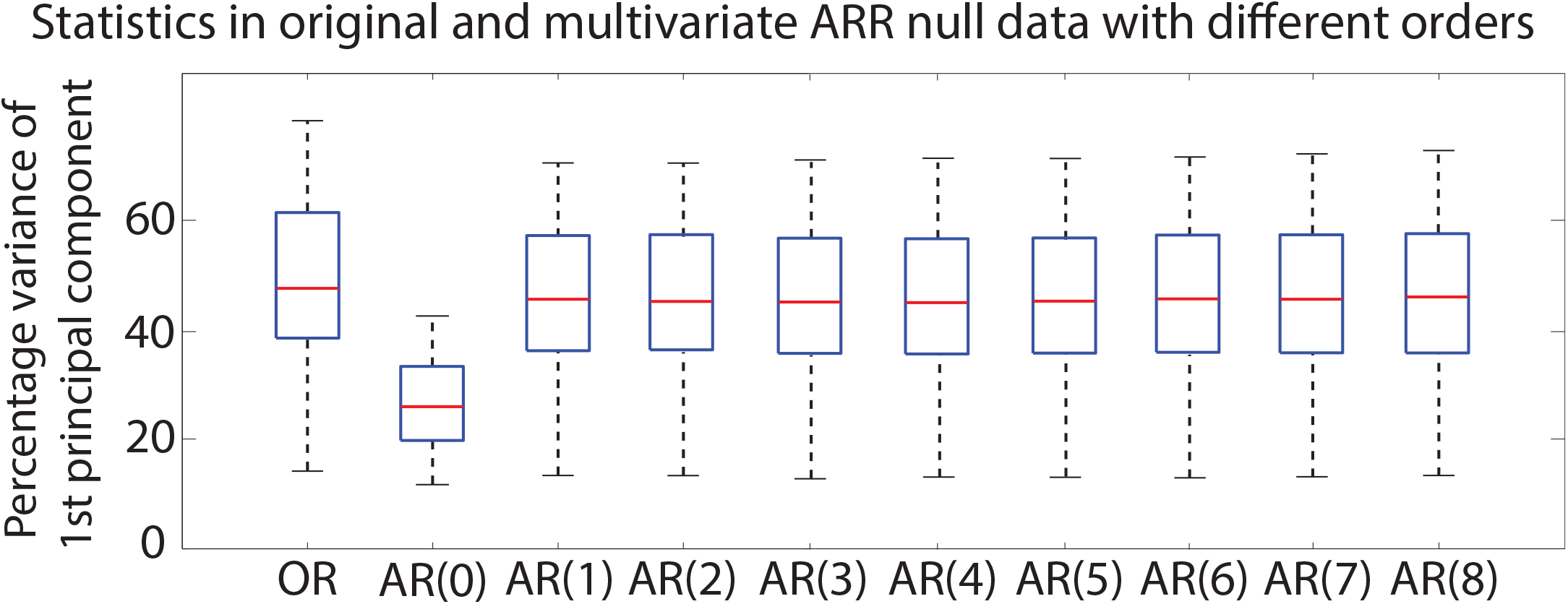
Impact of AR model order *p*. AR model of order 0 (static model) does not capture fluctuations in coherent brain dynamics (as measured by percentage variance of first principal component) present in the original (OR) data. AR models of orders 1 to 8 capture similar fluctuations in coherent brain dynamics.

### S4. Impact of temporal filtering and mean grayordinate regression (MGR)

**Figure S4:**
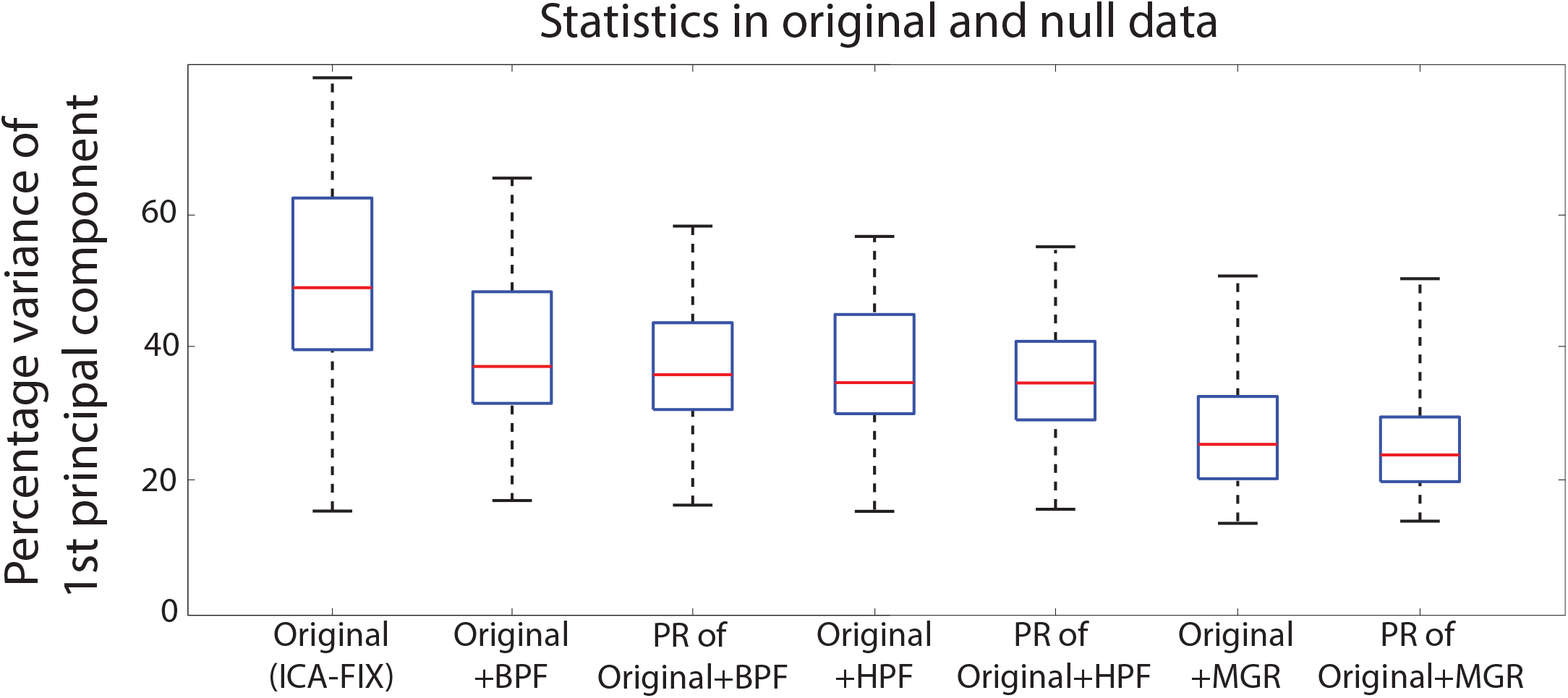
Impact of bandpass filtering (BPF), highpass filtering (HPF) and mean grayordinate regression (MGR). (Left) ICA-FIX original data. From left to right: bandpass filtered original data (0.01-0.1Hz) and corresponding phase randomized surrogates; highpass filtered original data (≥0.0167 Hz following the rule of thumb provided in Leonardi and Van De Ville (2015)); original data with mean grayordinate regression (MGR) and corresponding phase randomized surrogates. BPF, HPF and MGR reduce the percentage variance explained by the first principal component and the statistical difference between original and corresponding surrogates is also lowered.

### S5. Comparing Stationary Linear Gaussian Model and Hidden Markov Model

**Figure S5:**
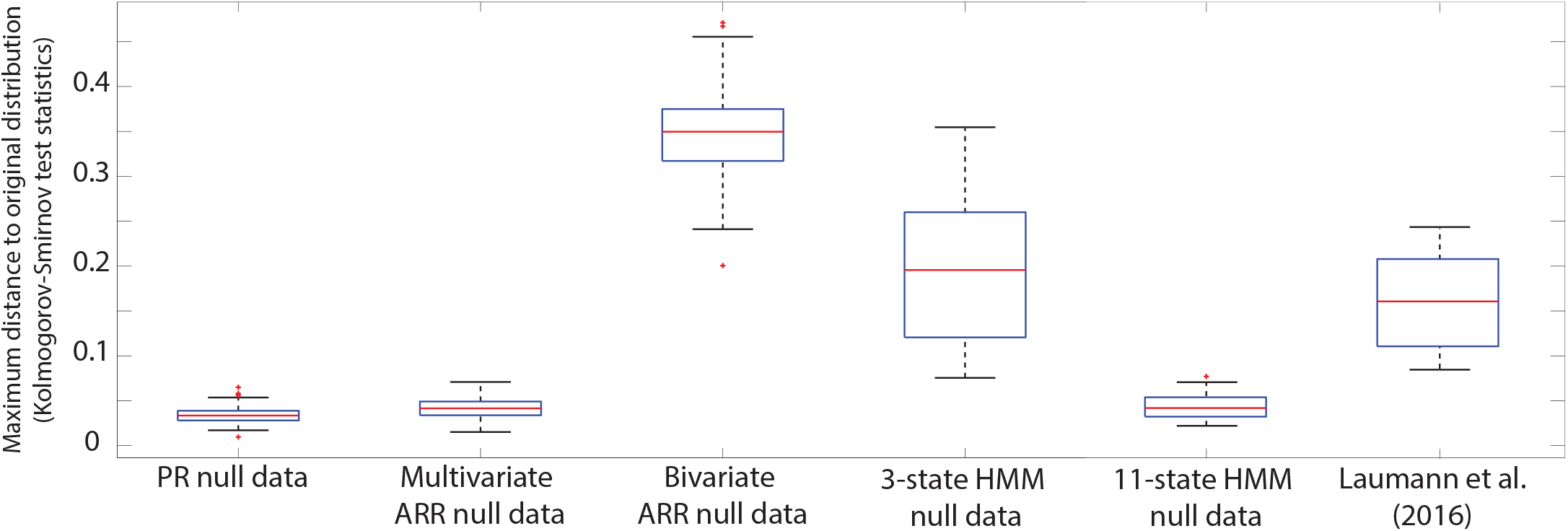
Distribution of Kolmogorov-Smirnov (KS) statistics between the functional connectivity dynamics (FCD) matrices of the original data and 100 null data from PR, multivariate ARR, bivariate ARR, 3-state hidden Markov model (HMM), 11-state HMM, and Laumann et al. (2016). A smaller KS statistic implies a better match between original data and null data as the KS statistics measures the maximum distance between two distributions.

### S6. FCD matrices computed with other surrogates

**Figure S6:**
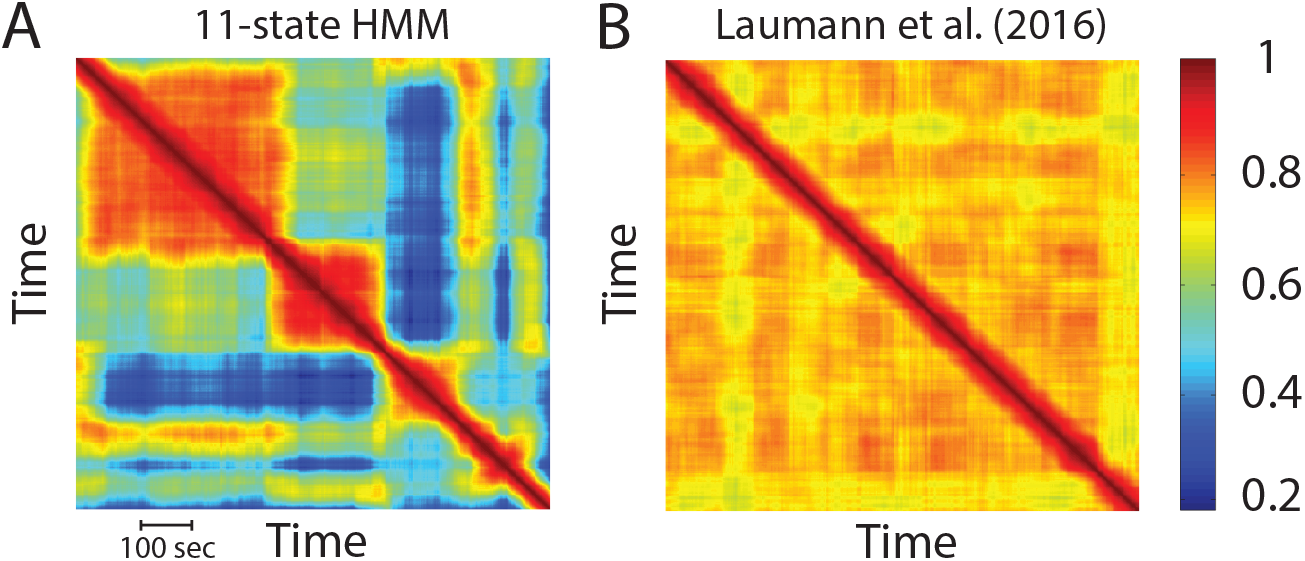
FCD matrices of (A) 11-state hidden Markov model (HMM) and (B) Laumann et al. (2016)

## Footnotes

1 Given recent interests in single subject analyses, there has been increasing amount of data collected for individual subjects, such as over 80 sessions of fMRI data for Russ Poldrack (Laumann et al., 2015; Poldrack et al., 2015; Braga and Buckner, 2017; Gordon et al., 2017). It is unclear to us whether different fMRI sessions of the same person at *different* times can be considered realizations of the *same* random process given obvious state differences, e.g., Russ was unfed/uncaffeinated on Tuesday and fed/caffeinated on Thursday. Ideally, we would like Russ to be scanned across multiple parallel sessions at the same time, which is obviously impossible. Furthermore, even if we were to treat the different sessions as realizations of the same random process, we would still face a power issue. For example, using a standard brain parcellation with 114 ROIs (Section 6), 80 fMRI sessions would not be sufficient to estimate a positive-definite (ensemble) covariance matrix for each TR (without additional regularization).

2 More generally, WSS processes encompass a wide range of signals because WSS can be the result of “interesting” dynamics being washed out across realizations. Indeed, given a stationary correlation time series, one can reverse engineer a bivariate random process exhibiting the correlation time series (like the HMM toy brain). This is not true for deterministic correlation time series, which is inevitably a result of non-stationary processes (assuming correlation is not a constant).

3 One could also modify the HMM so that the toy brain will be WSS, but exhibiting real dynamic spontaneous activity *and* dynamic FC. tual issues raised in this section, and dive into the statistical testing of non-stationarity when only a single realization of the random process is available.

4 The mean can always be added back to the null data hence there is no loss of generality to assume the original time courses were demeaned.

5 Σ might not be a diagonal matrix, i.e., the noise does not need to be independent across brain regions.

6 In statistical parlance, AR processes are dense in the class of linear Gaussian processes.

7 In reality, the procedure is slightly more complicated because the DFT of a real-valued time course (e.g., fMRI) exhibits Hermitian symmetry, i.e., the Fourier coefficient at frequency *k* is equal to the complex conjugate of the Fourier coefficient at frequency *T k*. As such, the random phases for half the frequencies determine the other half. See Appendix A3 for more details.

8 Values were obtained from Movement RelativeRMS.txt provided by HCP.

9 There were 1999 (and not 2000) null datasets because the real data is also counted as a dataset when computing *p* values, so that a *p* value of 0 is impossible.

10 Since there were 114 ROIs, each null dataset generated 6441 (= 114(114 1)*/*2) null values, which were pooled across the cortex into a single, highly-resolved null distribution (Zalesky et al., 2014). In other words, for a given participant, all the ROI pairs shared the same null distribution.

11 Following popular approaches (Allen et al., 2014; Wang et al., 2016), SWC of the representative participant were clustered into three states using k-means. Probability of transitioning between states were estimated based on the state assignment by k-means. Timepoints assigned to the same state was utilized to estimate the mean and covariance matrix of a multivariate Gaussian distribution using maximum likelihood. The estimated model parameters were then used to generate null data. It is worth noting that the 3-state HMM has roughly the same number of parameters as a first order multivariate AR model.

12 This is possible because rejection of the null hypothesis only implies the existence of information beyond the stationary, linear, Gaussian (SLG) model, and does not preclude the SLG model from explaining the original data better than an alternative model. For example, suppose 40% variance of the original data is uniquely explained by the SLG model, 10% variance is uniquely explained by an alternative model, and 50% variance is jointly explained by the SLG and alternative models. In this case, the alternative model explains some variance not explained by the SLG model. Therefore, using a metric derived from the alternative model can result in statistical rejection given enough statistical power. However, the SLG model is still better than the alternative model because it explains 90% of the data compared with 60% for the alternative model.

## References

Allen, E. A., E. Damaraju, S. M. Plis, E. B. Erhardt, T. Eichele, and V. D. Calhoun 2014. Tracking whole-brain connectivity dynamics in the resting state. Cerebral cortex, 24(3):663–676.

Anderson, J. S. 2008. Origin of synchronized low-frequency blood oxygen level–dependent fluctuations in the primary visual cortex. American Journal of Neuroradiology, 29(9):1722–1729.

Barttfeld, P., L. Uhrig, J. D. Sitt, M. Sigman, B. Jarraya, and S. Dehaene 2015. Signature of consciousness in the dynamics of resting-state brain activity. Proceedings of the National Academy of Sciences, 112(3):887–892.

Baum, L. E. and T. Petrie 1966. Statistical inference for probabilistic functions of finite state markov chains. The annals of mathematical statistics, 37(6):1554–1563.

Biswal, B., F. Z. Yetkin, V. M. Haughton, and J. S. Hyde 1995. Functional connectivity in the motor cortex of resting human brain using echo-planar mri. Magnetic resonance in medicine, 34:537–541.

Bollerslev, T. 1986. Generalized autoregressive conditional heteroskedasticity. Journal of econometrics, 31(3):307–327.

Braga, R. M. and R. L. Buckner 2017. Parallel interdigitated distributed networks within the individual estimated by intrinsic functional connectivity. Neuron, 95(2):457–471.

Breakspear, M., M. J. Brammer, E. T. Bullmore, P. Das, and L. M. Williams 2004. Spatiotemporal wavelet resampling for functional neuroimaging data. Human brain mapping, 23(1):1–25.

Buckner, R. L., F. M. Krienen, and B. T. Yeo 2013. Opportunities and limitations of intrinsic functional connectivity mri. Nature neuroscience, 16(7):832–837.

Buckner, R. L., J. Sepulcre, T. Talukdar, F. M. Krienen, H. Liu, T. Hedden, J. R. Andrews-Hanna, R. A. Sperling, and K. A. Johnson 2009. Cortical hubs revealed by intrinsic functional connectivity: mapping, assessment of stability, and relation to alzheimer’s disease. The Journal of neuroscience: the official journal of the Society for Neuroscience, 29:1860–1873.

Burgess, G. C., S. Kandala, D. Nolan, T. O. Laumann, J. D. Power, B. Adeyemo, M. P. Harms, S. E. Petersen, and D. M. Barch 2016. Evaluation of denoising strategies to address motion-correlated artifacts in resting-state functional magnetic resonance imaging data from the human connectome project. Brain connectivity, 6(9):669–680.

Calhoun, V. D., R. Miller, G. Pearlson, and T. Adalı 2014. The chronnectome: time-varying connectivity networks as the next frontier in fmri data discovery. Neuron, 84(2):262–274.

Casti, J. L. 1986. Linear dynamical systems. Academic Press Professional, Inc.

Chai, B., D. Walther, D. Beck, and L. Fei-Fei 2009. Exploring functional connectivities of the human brain using multivariate information analysis. In Advances in neural information processing systems, Pp. 270–278.

Chang, C. and G. H. Glover 2010. Time–frequency dynamics of resting-state brain connectivity measured with fmri. Neuroimage, 50(1):81–98.

Chang, C., D. A. Leopold, M. L. Schölvinck, H. Mandelkow, D. Picchioni, X. Liu, F. Q. Ye, J. N. Turchi, and J. H. Duyn 2016. Tracking brain arousal fluctuations with fmri. Proceedings of the National Academy of Sciences of the United States of America, 113(16):4518–4523.

Choe, A. S., M. B. Nebel, A. D. Barber, J. R. Cohen, Y. Xu, J. J. Pekar, B. Caffo, and M. A. Lindquist 2017. Comparing test-retest reliability of dynamic functional connectivity methods. NeuroImage.

Churchill, N. W., R. Spring, C. Grady, B. Cimprich, M. K. Askren, P. A. Reuter-Lorenz, M. S. Jung, S. Peltier, S. C. Strother, and M. G. Berman 2016. The suppression of scale-free fmri brain dynamics across three different sources of effort: aging, task novelty and task difficulty. Scientific reports, 6:30895.

Ciuciu, P., P. Abry, and B. J. He 2014. Interplay between functional connectivity and scale-free dynamics in intrinsic fmri networks. Neuroimage, 95:248–263.

Ciuciu, P., G. Varoquaux, P. Abry, S. Sadaghiani, and A. Kleinschmidt 2012. Scale-free and multifractal time dynamics of fmri signals during rest and task. Frontiers in physiology, 3.

Damaraju, E., E. Allen, A. Belger, J. Ford, S. McEwen, D. Mathalon, B. Mueller, G. Pearlson, S. Potkin, A. Preda, et al. 2014. Dynamic functional connectivity analysis reveals transient states of dysconnectivity in schizophrenia. NeuroImage: Clinical, 5:298–308.

Damoiseaux, J., S. Rombouts, F. Barkhof, P. Scheltens, C. Stam, S. M. Smith, and C. Beckmann 2006. Consistent resting-state networks across healthy subjects. Proceedings of the national academy of sciences, 103(37):13848–13853.

Deco, G., V. K. Jirsa, and A. R. McIntosh 2011. Emerging concepts for the dynamical organization of resting-state activity in the brain. Nature Reviews Neuroscience, 12(1):43–56.

Deco, G., V. K. Jirsa, P. A. Robinson, M. Breakspear, and K. Friston 2008. The dynamic brain: from spiking neurons to neural masses and cortical fields. PLoS Comput Biol, 4(8):e1000092.

Dosenbach, N. U., D. A. Fair, F. M. Miezin, A. L. Cohen, K. K. Wenger, R. A. Dosenbach, M. D. Fox, A. Z. Snyder, J. L. Vincent, M. E. Raichle, et al. 2007. Distinct brain networks for adaptive and stable task control in humans. Proceedings of the National Academy of Sciences, 104(26):11073–11078.

Du, Y., G. D. Pearlson, Q. Yu, H. He, D. Lin, J. Sui, L. Wu, and V. D. Calhoun 2016. Interaction among subsystems within default mode network diminished in schizophrenia patients: A dynamic connectivity approach. Schizophrenia research, 170(1):55–65.

Efron, B. and R. Tibshirani 1986. Bootstrap methods for standard errors, confidence intervals, and other measures of statistical accuracy. Statistical science, Pp. 54–75.

Eke, A., P. Herman, L. Kocsis, and L. Kozak 2002. Fractal characterization of complexity in temporal physiological signals. Physiological measurement, 23(1):R1.

El-Shaarawi, A. H. and W. W. Piegorsch 2013. Encyclopedia of environmetrics.

Fox, M. D., A. Z. Snyder, J. L. Vincent, and M. E. Raichle 2007. Intrinsic fluctuations within cortical systems account for intertrial variability in human behavior. Neuron, 56(1):171–184.

Fox, M. D., A. Z. Snyder, J. M. Zacks, and M. E. Raichle 2006. Coherent spontaneous activity accounts for trial-to-trial variability in human evoked brain responses. Nature neuroscience, 9(1):23–25.

Fransson, P. and G. Marrelec 2008. The precuneus/posterior cingulate cortex plays a pivotal role in the default mode network: Evidence from a partial correlation network analysis. Neuroimage, 42(3):1178–1184.

Freeman, W. J. and J. Zhai 2009. Simulated power spectral density (psd) of background electrocorticogram (ecog). Cognitive neurodynamics, 3(1):97–103.

Friston, K. 2009. Causal modelling and brain connectivity in functional magnetic resonance imaging. PLoS biol, 7(2):e1000033.

Friston, K. J. 2011. Functional and effective connectivity: a review. Brain connectivity, 1(1):13–36.

Friston, K. J., L. Harrison, and W. Penny 2003. Dynamic causal modelling. Neuroimage, 19(4):1273–1302.

Gajic, Z. 2003. Linear dynamic systems and signals. Prentice Hall – Pearson Education.

Glasser, M. F., S. N. Sotiropoulos, J. A. Wilson, T. S. Coalson, B. Fischl, J. L. Andersson, J. Xu, S. Jbabdi, M. Webster, J. R. Polimeni, et al. 2013. The minimal preprocessing pipelines for the human connectome project. Neuroimage, 80:105–124.

Gordon, E. M., T. O. Laumann, A. W. Gilmore, D. J. Newbold, D. J. Greene, J. J. Berg, M. Ortega, C. Hoyt-Drazen, C. Gratton, H. Sun, et al. 2017. Precision functional mapping of individual human brains. Neuron.

Grandjean, J., M. G. Preti, T. A. Bolton, M. Buerge, E. Seifritz, C. R. Pryce, D. Van De Ville, and M. Rudin 2017. Dynamic reorganization of intrinsic functional networks in the mouse brain. NeuroImage, 152:497–508.

Greicius, M. D., B. Krasnow, A. L. Reiss, and V. Menon 2003. Functional connectivity in the resting brain: a network analysis of the default mode hypothesis. Proceedings of the National Academy of Sciences, 100(1):253–258.

Griffa, A., B. Ricaud, K. Benzi, X. Bresson, A. Daducci, P. Vandergheynst, J.-P. Thiran, and P. Hagmann 2017. Transient networks of spatio-temporal connectivity map communication pathways in brain functional systems. NeuroImage – In Press.

Handwerker, D. A., V. Roopchansingh, J. Gonzalez-Castillo, and P. A. Bandettini 2012. Periodic changes in fmri connectivity. Neuroimage, 63(3):1712–1719.

Hansen, E. C., D. Battaglia, A. Spiegler, G. Deco, and V. K. Jirsa 2015. Functional connectivity dynamics: modeling the switching behavior of the resting state. Neuroimage, 105:525–535.

Hathout, G. M., R. K. Gopi, P. Bandettini, and S. S. Gambhir 1999. The lag of cerebral hemodynamics with rapidly alternating periodic stimulation: modeling for functional mri. Magnetic resonance imaging, 17(1):9–20.

He, B. J. 2011. Scale-free properties of the functional magnetic resonance imaging signal during rest and task. Journal of Neuroscience, 31(39):13786–13795.

He, B. J. 2014. Scale-free brain activity: past, present, and future. Trends in cognitive sciences, 18(9):480–487.

He, B. J. and M. E. Raichle 2009. The fmri signal, slow cortical potential and consciousness. Trends in cognitive sciences, 13(7):302–309.

He, B. J., J. M. Zempel, A. Z. Snyder, and M. E. Raichle 2010. The temporal structures and functional significance of scale-free brain activity. Neuron, 66(3):353–369.

Hindriks, R., M. H. Adhikari, Y. Murayama, M. Ganzetti, D. Mantini, N. K. Logothetis, and G. Deco 2016. Can sliding-window correlations reveal dynamic functional connectivity in resting-state fmri? Neuroimage, 127:242–256.

Hodgkin, A. L. and A. F. Huxley 1952. A quantitative description of membrane current and its application to conduction and excitation in nerve. The Journal of physiology, 117(4):500.

Hutchison, R. M., T. Womelsdorf, E. A. Allen, P. A. Bandettini, V. D. Calhoun, M. Corbetta, S. Della Penna, J. H. Duyn, G. H. Glover, J. Gonzalez-Castillo, et al. 2013a. Dynamic functional connectivity: promise, issues, and interpretations. Neuroimage, 80:360–378.

Hutchison, R. M., T. Womelsdorf, J. S. Gati, S. Everling, and R. S. Menon 2013b. Resting-state networks show dynamic functional connectivity in awake humans and anesthetized macaques. Human brain mapping, 34(9):2154–2177.

Karahanoğlu, F. I. and D. Van De Ville 2015. Transient brain activity disentangles fmri resting-state dynamics in terms of spatially and temporally overlapping networks. Nature communications, 6.

Khintchine, A. 1934. Korrelationstheorie der stationaryren stochastischen prozesse. Mathematische Annalen, 109(1):604–615.

Kragel, P. A., A. R. Knodt, A. R. Hariri, and K. S. LaBar 2016. Decoding spontaneous emotional states in the human brain. PLoS biology, 14(9):e2000106.

Krienen, F. M., B. T. Yeo, T. Ge, R. L. Buckner, and C. C. Sherwood 2016. Transcriptional profiles of supragranular-enriched genes associate with corticocortical network architecture in the human brain. Proceedings of the National Academy of Sciences, 113(4):E469–E478.

Kucyi, A., M. J. Hove, M. Esterman, R. M. Hutchison, and E. M. Valera 2016. Dynamic brain network correlates of spontaneous fluctuations in attention. Cerebral Cortex, 27(3):1831–1840.

Laumann, T. O., E. M. Gordon, B. Adeyemo, A. Z. Snyder, S. J. Joo, M.-Y. Chen, A. W. Gilmore, K. B. McDermott, S. M. Nelson, N. U. F. Dosenbach, B. L. Schlaggar, J. A. Mumford, R. A. Poldrack, and S. E. Petersen 2015. Functional system and areal organization of a highly sampled individual human brain. Neuron, 87(3):657–670.

Laumann, T. O., A. Z. Snyder, A. Mitra, E. M. Gordon, C. Gratton, B. Adeyemo, A. W. Gilmore, S. M. Nelson, J. J. Berg, D. J. Greene, et al. 2016. On the stability of bold fmri correlations. Cerebral Cortex, Pp. 1–14.

Leonardi, N., W. R. Shirer, M. D. Greicius, and D. Van De Ville 2014. Disentangling dynamic networks: Separated and joint expressions of functional connectivity patterns in time. Human brain mapping, 35(12):5984–5995.

Leonardi, N. and D. Van De Ville 2015. On spurious and real fluctuations of dynamic functional connectivity during rest. Neuroimage, 104:430–436.

Liégeois, R. 2015. Dynamical modelling from resting-state brain imaging. PhD thesis, University of Liège, Liège, Belgium.

Liégeois, R., B. Mishra, M. Zorzi, and R. Sepulchre 2015. Sparse plus low-rank autoregressive identification in neuroimaging time series. In Proc. 54th IEEE Conf. Decision and Control (CDC), Pp. 3965–3970.

Liégeois, R., E. Ziegler, C. Phillips, P. Geurts, F. Gómez, M. A. Bahri, B. T. T. Yeo, A. Soddu, A. Vanhaudenhuyse, S. Laureys, and R. Sepulchre 2016. Cerebral functional connectivity periodically (de)synchronizes with anatomical constraints. Brain structure and function, 221:2985–2997.

Lindquist, M. A., Y. Xu, M. B. Nebel, and B. S. Caffo 2014. Evaluating dynamic bivariate correlations in resting-state fmri: A comparison study and a new approach. Neuroimage, 101:531–546.

Lutkepohl, H. 2005. New introduction to multiple time series analysis. Econometric theory, 22(5):961–967.

Maiwald, T., E. Mammen, S. Nandi, and J. Timmer 2008. Surrogate Data — A Qualitative and Quantitative Analysis, Pp. 41–74. Berlin, Heidelberg: Springer Berlin Heidelberg.

Majeed, W., M. Magnuson, W. Hasenkamp, H. Schwarb, E. H. Schumacher, L. Barsalou, and S. D. Keilholz 2011. Spatiotemporal dynamics of low frequency bold fluctuations in rats and humans. Neuroimage, 54(2):1140–1150.

Margulies, D. S., S. S. Ghosh, A. Goulas, M. Falkiewicz, J. M. Huntenburg, G. Langs, G. Bezgin, S. B. Eickhoff, F. X. Castellanos, M. Petrides, et al. 2016. Situating the default-mode network along a principal gradient of macroscale cortical organization. Proceedings of the National Academy of Sciences, 113(44):12574–12579.

Matsui, T., T. Murakami, and K. Ohki 2016. Transient neuronal coactivations embedded in globally propagating waves underlie resting-state functional connectivity. Proceedings of the National Academy of Sciences, 113(23):6556–6561.

Miller, K. J., L. B. Sorensen, J. G. Ojemann, and M. Den Nijs 2009. Power-law scaling in the brain surface electric potential. PLoS computational biology, 5(12):e1000609.

Miller, R. L., M. Yaesoubi, J. A. Turner, D. Mathalon, A. Preda, G. Pearlson, T. Adali, and V. D. Calhoun 2016. Higher dimensional meta-state analysis reveals reduced resting fmri connectivity dynamism in schizophrenia patients. PloS one, 11(3):e0149849.

Milstein, J., F. Mormann, I. Fried, and C. Koch 2009. Neuronal shot noise and brownian 1/f2 behavior in the local field potential. PloS one, 4(2):e4338.

Mitra, A. and M. E. Raichle 2016. How networks communicate: propagation patterns in spontaneous brain activity. Phil. Trans. R. Soc. B, 371(1705):20150546.

Mitra, A., A. Z. Snyder, C. D. Hacker, and M. E. Raichle 2014. Lag structure in resting-state fmri. Journal of neurophysiology, 111(11):2374–2391.

Mohajerani, M. H., A. W. Chan, M. Mohsenvand, J. LeDue, R. Liu, D. A. McVea, J. D. Boyd, Y. T. Wang, M. Reimers, and T. H. Murphy 2013. Spontaneous cortical activity alternates between motifs defined by regional axonal projections. Nature neuroscience, 16(10):1426–1435.

Nason, G. 2013. A test for second-order stationarity and approximate confidence intervals for localized autocovariances for locally stationary time series. Journal of the Royal Statistical Society: Series B (Statistical Methodology), 75(5):879–904.

Nomi, J. S., S. G. Vij, D. R. Dajani, R. Steimke, E. Damaraju, S. Rachakonda, V. D. Calhoun, and L. Q. Uddin 2017. Chronnectomic patterns and neural flexibility underlie executive function. NeuroImage, 147:861–871.

Oppenheim, A. and A. S. Willsky 1997. Signals and Systems.

Osborne, A., A. Kirwan, A. Provenzale, and L. Bergamasco 1986. A search for chaotic behavior in large and mesoscale motions in the pacific ocean. Physica D: Nonlinear Phenomena, 23(1):75–83.

Papoulis, A. 2002. Probability, Random Variables and Stochastic Processes. Mc Graw Hill.

Pfaff, B. 2008. Analysis of integrated and cointegrated time series with R. Springer Science & Business Media.

Poldrack, R. A., T. O. Laumann, O. Koyejo, B. Gregory, A. Hover, M.-Y. Chen, K. J. Gorgolewski, J. Luci, S. J. Joo, R. L. Boyd, et al 2015. Long-term neural and physiological phenotyping of a single human. Nature communications, 6:8885.

Pollock, D. 2011. Econometric Theory. University of Leicester.

Power, J. D., A. L. Cohen, S. M. Nelson, G. S. Wig, K. A. Barnes, J. A. Church, A. C. Vogel, T. O. Laumann, F. M. Miezin, B. L. Schlaggar, and S. E. Petersen 2011. Functional network organization of the human brain. Neuron, 72(4):665–678.

Power, J. D., M. Plitt, T. O. Laumann, and A. Martin 2017. Sources and implications of whole-brain fmri signals in humans. Neuroimage, 146:609–625.

Preti, M. G., T. A. Bolton, and D. Van De Ville 2016. The dynamic functional connectome: State-of-the-art and perspectives. NeuroImage – In Press.

Prichard, D. and J. Theiler 1994. Generating surrogate data for time series with several simultaneously measured variables. Physical Review Letters, 73(7):951–954.

Prince, S. J. 2012. Computer vision: models, learning, and inference. Cambridge University Press.

Raatikainen, V., N. Huotari, V. Korhonen, A. Rasila, J. Kananen, L. Raitamaa, T. Keinänen, J. Kantola, O. Tervonen, and V. Kiviniemi 2017. Combined spatiotemporal ica (stica) for continuous and dynamic lag structure analysis of mreg data. NeuroImage.

Rabiner, L. R. 1989. A tutorial on hidden markov models and selected applications in speech recognition. Proceedings of the IEEE, 77(2):257–286.

Rashid, B., E. Damaraju, G. D. Pearlson, and V. D. Calhoun 2014. Dynamic connectivity states estimated from resting fmri identify differences among schizophrenia, bipolar disorder, and healthy control subjects. Frontiers in human neuroscience, 8:897.

Rogers, B. P., S. B. Katwal, V. L. Morgan, C. L. Asplund, and J. C. Gore 2010. Functional mri and multivariate autoregressive models. Magnetic resonance imaging, 28(8):1058–1065.

Sakoğlu, Ü., G. D. Pearlson, K. A. Kiehl, Y. M. Wang, A. M. Michael, and V. D. Calhoun 2010. A method for evaluating dynamic functional network connectivity and task-modulation: application to schizophrenia. Magnetic Resonance Materials in Physics, Biology and Medicine, 23(5-6):351–366.

Schaefer, A., R. Kong, E. M. Gordon, T. O. Laumann, X.-N. Zuo, A. Holmes, S. B. Eickhoff, and B. T. Yeo 2017. Local-global parcellation of the human cerebral cortex from intrinsic functional connectivity mri. Cerebral Cortex, In Press, P. 135632.

Schreiber, T. and A. Schmitz 2000. Surrogate time series. Physica D: Nonlinear Phenomena, 142(3):346–382.

Shine, J. M., O. Koyejo, P. T. Bell, K. J. Gorgolewski, M. Gilat, and R. A. Poldrack 2015. Estimation of dynamic functional connectivity using multiplication of temporal derivatives. NeuroImage, 122:399–407.

Shine, J. M., O. Koyejo, and R. A. Poldrack 2016. Temporal metastates are associated with differential patterns of time-resolved connectivity, network topology, and attention. Proceedings of the National Academy of Sciences, 113(35):9888–9891.

Siegel, J. S., A. Mitra, T. O. Laumann, B. A. Seitzman, M. Raichle, M. Corbetta, and A. Z. Snyder 2016. Data quality influences observed links between functional connectivity and behavior. Cerebral Cortex – In Press.

Smith, S. M., C. F. Beckmann, J. Andersson, E. J. Auerbach, J. Bijsterbosch, G. Douaud, E. Duff, D. A. Feinberg, L. Griffanti, M. P. Harms, M. Kelly, T. Laumann, K. L. Miller, S. Moeller, S. Petersen, J. Power, G. Salimi-Khorshidi, A. Z. Snyder, A. T. Vu, M. W. Woolrich, J. Xu, E. Yacoub, K. Ugurbil, D. C. V. Essen, and M. F. Glasser 2013a. Resting-state fmri in the human connectome project. NeuroImage, 80:144–168.

Smith, S. M., K. L. Miller, S. Moeller, J. Xu, E. J. Auerbach, M. W. Woolrich, C. F. Beckmann, M. Jenkinson, J. Andersson, M. F. Glasser, et al. 2012. Temporally-independent functional modes of spontaneous brain activity. Proceedings of the National Academy of Sciences, 109(8):3131–3136.

Smith, S. M., D. Vidaurre, C. F. Beckmann, M. F. Glasser, M. Jenkinson, K. L. Miller, T. E. Nichols, E. C. Robinson, G. Salimi-Khorshidi, M. W. Woolrich, et al. 2013b. Functional connectomics from resting-state fmri. Trends in cognitive sciences, 17(12):666–682.

Spreng, R. N., J. Sepulcre, G. R. Turner, W. D. Stevens, and D. L. Schacter 2013. Intrinsic architecture underlying the relations among the default, dorsal attention, and frontoparietal control networks of the human brain. Journal of cognitive neuroscience, 25(1):74–86.

Stephan, K. E., L. Kasper, L. M. Harrison, J. Daunizeau, H. E. den Ouden, M. Breakspear, and K. J. Friston 2008. Nonlinear dynamic causal models for fmri. Neuroimage, 42(2):649–662.

Stoica, P. and R. L. Moses 2005. Spectral analysis of signals. Pearson/Prentice Hall Upper Saddle River, NJ.

Stoica, P. and A. Nehorai 1987. On stability and root location of linear prediction models. IEEE Transactions on Acoustics, Speech, and Signal Processing, 35(4):582–584.

Stroh, A., H. Adelsberger, A. Groh, C. RÜhlmann, S. Fischer, A. Schierloh, K. Deisseroth, and A. Konnerth 2013. Making waves: initiation and propagation of corticothalamic ca 2+ waves in vivo. Neuron, 77(6):1136–1150.

Su, J., H. Shen, L.-L. Zeng, J. Qin, Z. Liu, and D. Hu 2016. Heredity characteristics of schizophrenia shown by dynamic functional connectivity analysis of resting-state functional mri scans of unaffected siblings. NeuroReport, 27(11):843–848.

Tedeschi, W., H.-P. MÜller, D. de Araujo, A. Santos, U. Neves, S. Ernè, and O. Baffa 2005. Generalized mutual information tests applied to fmri analysis. Physica A: Statistical Mechanics and its Applications, 352(2):629–644.

Theiler, J., S. Eubank, A. Longtin, B. Galdrikian, and J. D. Farmer 1992. Testing for nonlinearity in time series: the method of surrogate data. Physica D: Nonlinear Phenomena, 58(1):77–94.

Ting, C.-M., A.-K. Seghouane, and S.-H. Salleh 2016. Estimation of high-dimensional connectivity in fmri data via subspace autoregressive models. In Statistical Signal Processing Workshop (SSP), 2016 IEEE, Pp. 1–5. IEEE.

Tsai, A., J. W. Fisher III, C. Wible, W. M. Wells III, J. Kim, and A. S. Willsky 1999. Analysis of functional mri data using mutual information. In International Conference on Medical Image Computing and Computer-Assisted Intervention, Pp. 473–480.

Tucker, M., P. G. Challenor, and D. Carter 1984. Numerical simulation of a random sea: a common error and its effect upon wave group statistics. Applied ocean research, 6(2):118–122.

Valdes, P., J. C. Jiménez, J. Riera, R. Biscay, and T. Ozaki 1999. Nonlinear eeg analysis based on a neural mass model. Biological cybernetics, 81(5):415–424.

Van De Ville, D., T. Blu, and M. Unser 2004. Integrated wavelet processing and spatial statistical testing of fmri data. NeuroImage, 23(4):1472–1485.

Van de Ville, D., J. Britz, and C. M. Michel 2010. Eeg microstate sequences in healthy humans at rest reveal scale-free dynamics. Proceedings of the National Academy of Sciences, 107(42):18179–18184.

Van Den Heuvel, M. P. and H. E. H. Pol 2010. Exploring the brain network: a review on resting-state fmri functional connectivity. European neuropsychopharmacology, 20(8):519–534.

Van Essen, D. C., S. M. Smith, D. M. Barch, T. E. Behrens, E. Yacoub, K. Ugurbil, W.-M. H. Consortium, et al. 2013. The wu-minn human connectome project: an overview. Neuroimage, 80:62–79.

Vincent, J. L., A. Z. Snyder, M. D. Fox, B. J. Shannon, J. R. Andrews, M. E. Raichle, and R. L. Buckner 2006. Coherent spontaneous activity identifies a hippocampal-parietal memory network. Journal of neurophysiology, 96(6):3517–3531.

Walker, G. 1931. On periodicity in series of related terms. Proceedings of the Royal Society of London. Series A, Containing Papers of a Mathematical and Physical Character, 131(818):518–532.

Wang, C., J. L. Ong, A. Patanaik, J. Zhou, and M. W. Chee 2016. Spontaneous eyelid closures link vigilance fluctuation with fmri dynamic connectivity states. Proceedings of the National Academy of Sciences, 113(34):9653–9658.

Weisstein, E. 2016. ”cross-correlation theorem.” from mathworld–a wolfram web resource.

Wiener, N. 1930. Generalized harmonic analysis. Acta Mathematica, 55:117–258.

Yeo, B. T., F. M. Krienen, M. W. Chee, and R. L. Buckner 2014. Estimates of segregation and overlap of functional connectivity networks in the human cerebral cortex. Neuroimage, 88:212–227.

Yeo, B. T. T., F. M. Krienen, J. Sepulcre, M. R. Sabuncu, D. Lashkari, M. Hollinshead, J. L. Roffman, J. W. Smoller, L. Zöllei, J. R. Polimeni, B. Fischl, H. Liu, and R. L. Buckner 2011. The organization of the human cerebral cortex estimated by intrinsic functional connectivity. Journal of neurophysiology, 106:1125–1165.

Yeo, B. T. T., J. Tandi, and M. W. L. Chee 2015. Functional connectivity during rested wakefulness predicts vulnerability to sleep deprivation. NeuroImage, 111:147–158.

Yule, G. U. 1927. On a method of investigating periodicities in disturbed series, with special reference to wolfer’s sunspot numbers. Philosophical Transactions of the Royal Society of London. Series A, Containing Papers of a Mathematical or Physical Character, 226:267–298.

Zalesky, A. and M. Breakspear 2015. Towards a statistical test for functional connectivity dynamics. NeuroImage, 114:466–470.

Zalesky, A., A. Fornito, and E. T. Bullmore 2010. Network-based statistic: identifying differences in brain networks. Neuroimage, 53(4):1197–1207.

Zalesky, A., A. Fornito, L. Cocchi, L. L. Gollo, and M. Breakspear 2014. Time-resolved resting-state brain networks. Proceedings of the National Academy of Sciences, 111(28):10341–10346.

Zivot, E. and J. Wang 2006. Vector autoregressive models for multivariate time series. Modeling Financial Time Series with S-PLUS, Pp. 385–429.

